# TWIST: A diagnostic framework for representing tree water deficit dynamics in process-based forest models

**DOI:** 10.64898/2026.06.15.732331

**Authors:** Yanick Ziegler, Pia Labenski, Martin Thurner, Jan Krejza, Ladislav Šigut, Nadine K. Ruehr, Rüdiger Grote

## Abstract

Dendrometer-derived tree water deficit (TWD) contains physiologically rich information and is increasingly used to monitor tree drought stress, yet process-based forest models rarely include a directly comparable representation of TWD dynamics. Existing hydraulic models can represent internal water storage in detail, but their parameter demands limit broader application. Here, we introduce the Tree Water Imbalance and Storage Tracker (TWIST), a parsimonious and physiologically interpretable framework that derives volume-based TWD dynamics. The module is driven by transpiration and relative soil water content and uses three empirical parameters to control transpiration-driven internal water depletion, deficit refilling, and additional soil-water uptake limitation. It also derives relative tree water content (RWC_tree_) from the simulated deficit and an estimate of the available internal water pool. We tested TWIST by coupling it to the process-based ecosystem model LandscapeDNDC. Parameters were optimized for 2018 and evaluated independently for 2019–2024 against normalized dendrometer-derived TWD at a Czech beech site. Simulated TWD trajectories broadly agreed with observed daily and seasonal dynamics, while RWC_tree_ translated them into a physiologically interpretable proxy for internal dehydration. TWIST demonstrated capability to reproduce key TWD drought-response patterns, including diurnal depletion–replenishment cycles, reduced nocturnal rehydration with declining soil moisture, and progressive deficit accumulation. By representing TWD and RWC_tree_ as diagnostic model outputs, TWIST makes dendrometer-derived drought-stress information more directly usable in forest models. It thereby provides a practical basis for linking tree-level drought-stress signals with stand-level simulations and, potentially, remotely sensed indicators of canopy water status.

## 1 Introduction

Internal tree water storage plays a crucial role in buffering imbalances between transpirational water loss and soil water uptake (Salomón et al., 2017; Preisler et al., 2022). While storage contributes substantially to daily transpiration (Čermák et al., 2007; Verbeeck et al., 2007; Betsch et al., 2011; Köcher et al., 2013; Preisler et al., 2022), it becomes particularly critical under drought, when soil water supply is limited (Klein et al., 2014; Klein et al., 2016; Preisler et al., 2019). Water released from tree tissues can prolong the duration of dehydration and delay the onset of hydraulic damage (Gleason et al., 2014). Even with fully closed stomata, residual water losses persist (Blackman et al., 2019; Duursma et al., 2019), and progressive dehydration can ultimately lead to hydraulic failure and cellular damage (Mantova et al., 2021; Mantova et al., 2022; McDowell et al., 2022). Consequently, internal water storage dynamics are fundamental for understanding the progression of drought stress and the persistence of long-term drought legacy effects (Müller and Bahn, 2022).

The relevance of such observations is amplified under ongoing climate change, which is increasing the frequency, duration, and intensity of drought events (Gazol et al., 2025; Chen et al., 2025). Such droughts already cause widespread growth limitations (Bose et al., 2025) and forest mortality across Europe (Senf et al., 2020) and worldwide (International Tree Mortality Network, 2025).

Continuous observations of tree water status are therefore increasingly important for detecting drought stress in forests (Novick et al., 2022; Restrepo-Acevedo et al., 2026). High-resolution dendrometers provide such observations by recording reversible stem shrinkage, from which tree water deficit (TWD) can be derived as an indicator of tree hydration status (Zweifel et al., 2001; Zweifel and Häsler, 2001). TWD carries physiologically rich information (De Swaef et al., 2015; Sevanto, 2025) due to its close relation to *ψ* (Dietrich et al., 2018; Ziegler et al., 2024) and its direct link to cell turgor pressure and growth processes (Zweifel et al., 2021a; Peters et al., 2023). Daily TWD fluctuations integrate atmospheric demand, soil water supply (Zweifel et al., 2005; Oberhuber et al., 2023), and species-specific water-use strategies (Sánchez-Costa et al., 2015; Peters et al., 2025), whereas unrecovered TWD over consecutive days indicates sustained drought stress and may reveal thresholds related to stomatal closure, turgor loss, and hydraulic impairment (Andriantelomanana et al., 2024; Ziegler et al., 2024; Peters et al., 2025). With the growing availability of long-term data from coordinated dendrometer networks, including the Swiss TreeNet (Zweifel et al., 2021b), the Czech DendroNetwork (https://dendronet.cz/), and the Global Dendrometer Network (https://globaldendro.org/), TWD is increasingly used to monitor tree drought stress. However, this growing observational capacity is not yet matched by a comparable representation of TWD dynamics in process-based forest models.

Process-based models are essential tools for estimating the effects of drought on forests. They allow scaling physiological processes from individual trees to stands, integrate multiple stressors over time, and can explore drought conditions beyond the observed range – providing insights that are difficult to obtain from measurements or statistical modeling alone. However, while current process-based schemes often capture drought effects on stomatal conductance, they rarely include variables directly comparable to dendrometer-derived TWD. This limits the use of dendrometer observations to evaluate modeled drought-stress dynamics and restricts the transfer of the physiological information contained in TWD into model frameworks. In addition, the effects of drought on tissue integrity, growth, and mortality remain only partially represented, as the underlying physiological mechanisms are insufficiently captured (Hartmann et al., 2018).

Mechanistic representations of drought stress often rely on tree water potential (*ψ*) or hydraulic impairment metrics. However, accurate *ψ* simulations remain challenging due to its non-linear, species-specific behavior and its sensitivity to soil texture – a highly variable site property. Drought-induced mortality is usually associated with percentage loss of conductivity (PLC) thresholds, as implemented in models such as ParFlow-TREES (Tai et al., 2018) or LPJ-GUESS (Meyer et al., 2025), but such approaches are difficult to constrain empirically due to species-specific vulnerability curves and methodological uncertainties (Cochard et al., 2013; Li et al., 2016; Feng et al., 2023).

Given these limitations, a stronger focus on tree water pools, which can be represented at a more integrative scale, may improve our ability to understand and anticipate drought impacts, as highlighted by Martinez-Vilalta et al. (2019). Despite increasing efforts to incorporate plant hydraulics into ecosystem frameworks, such explicit formulations of internal water storage and deficit dynamics remain rare. A meaningful representation of tree water status requires indicators that capture internal storage dynamics, respond consistently to dehydration, and can be directly compared with observations. TWD and relative tree water content (RWC_tree_) could meet these criteria.

While TWD provides an observation-linked measure of internal water deficit dynamics, RWC_tree_ complements it by expressing modeled deficits relative to available internal water storage. As the ratio of current to fully saturated water content, it reflects the physical consequences of cellular dehydration (Martinez-Vilalta et al., 2019) and has been linked to multiple drought-induced processes, including declines in stomatal and hydraulic conductance. Previous studies have also associated RWC with key physiological transitions such as turgor loss, metabolic impairment, and cellular damage (Bartlett et al., 2012; Lawlor and Cornic, 2002; Trueba et al., 2019; Sapes and Sala, 2021; Trifilò et al., 2023). Compared with *ψ*, RWC-based thresholds appear to show less inter-specific and organ-specific variability (Trueba et al., 2019; Trifilò et al., 2023), although uncertainty remains regarding their measurement and their relationship to hydraulic failure (Martinez-Vilalta et al., 2019). Nevertheless, RWC_tree_ can provide a physiologically interpretable proxy of relative internal hydration, with potential relevance for ecosystem-scale assessments of vegetation water content (Binks et al., 2024).

Representing TWD and RWC_tree_ in process-based forest models could create a direct interface between tree-level drought monitoring and model-based drought assessment. Several existing approaches can reproduce stem diameter variation, ranging from detailed mechanistic models (Peramaki et al., 2001; Peramaki et al., 2005; Steppe et al., 2006) to formulations designed primarily for interpreting dendrometer signals (Sevanto, 2001; Zweifel et al., 2005). In parallel, detailed hydraulic models such as SurEau and FETCH4 (Cochard et al., 2021; Ruffault et al., 2022; Missik et al., 2025), as well as hydraulically enhanced ecosystem models such as ED2-Hydro, ORCHIDEE-CAN-HYD, and FATES-HYDRO (Xu et al., 2016; Yao et al., 2022; Xu et al., 2023) already simulate tissue water storage and related hydraulic processes. These approaches provide important physiological detail, but they were generally not designed to yield a simple, diagnostically useful TWD state variable that can be constrained using dendrometer observations. A complementary need therefore remains for physiologically interpretable formulations that require few empirical parameters and make dendrometer-observable TWD dynamics usable in model simulations.

Therefore, in this study, we develop and test a tree water module, termed TWIST, that derives TWD dynamics from transpiration and relative soil water content and additionally estimates RWC_tree_ from modeled deficits and available internal water storage. TWIST is designed as a diagnostic add-on module that provides a parsimonious, physiologically interpretable representation of internal water deficit dynamics and allows model outputs to be compared with dendrometer-derived TWD. Specifically, we (i) present the module and illustrate its behavior under contrasting soil moisture conditions, (ii) integrate it within the process-based model LandscapeDNDC (Haas et al., 2013; Nadal-Sala et al., 2024) and evaluate it using multi-year dendrometer observations from a temperate beech forest site, and (iii) explore its response under an idealized rainfall-suppression stress test to examine boundary-case model behavior.

## 2 Materials and Methods

In this section, we first introduce the Tree Water Imbalance and Storage Tracker (TWIST) module, a lightweight water storage framework that simulates volume-based TWD and RWC_tree_ (Section 2.1), and then describe how it is coupled to the process-based ecosystem model LandscapeDNDC (Section 2.2). We next summarize the study site, observational data, normalization procedure, parameter calibration, and uncertainty analysis used for model evaluation against multi-year dendrometer observations (Sections 2.3–2.5). Finally, we introduce the “rainfall-suppression stress test”, designed to assess model behavior under imposed soil-water depletion (Section 2.6).

### 2.1 The TWIST Module

#### 2.1.1 Simulating tree water deficit dynamics

TWIST is a simple diagnostic mass-balance module that represents internal tree water storage and deficit as the cumulative imbalance between transpirational demand (*E*) and soil water uptake (*U*), modulated by soil moisture limitation and species-specific rehydration capacity (Fig. 1). It was developed as a lightweight, mechanistically inspired, and physiologically interpretable set of equations that can be coupled to process-based ecosystem models, without resolving explicit *ψ* gradients or tissue-specific hydraulic properties. An R implementation is provided for reproducibility (see Data and code availability).

**Figure 1:**
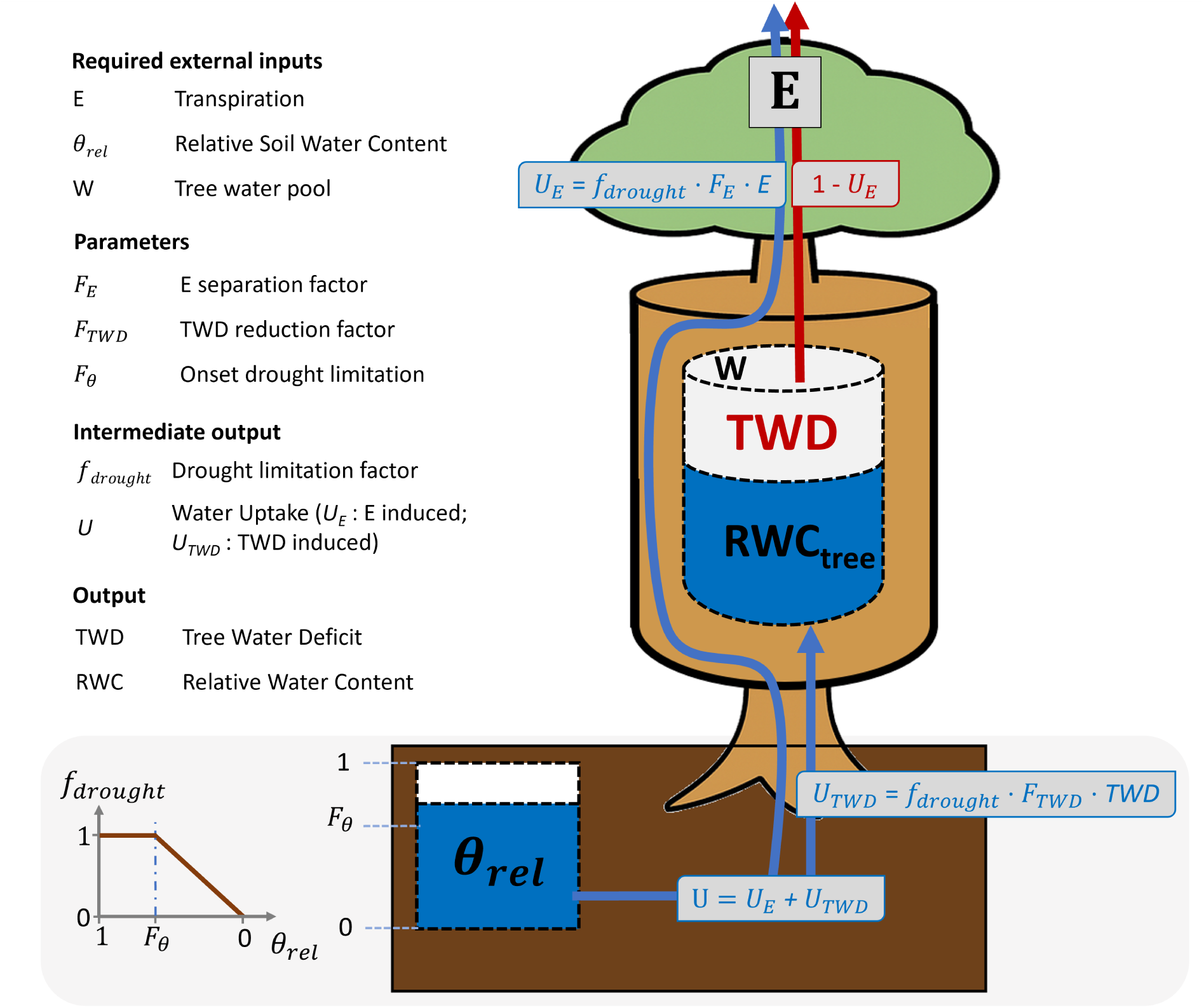
Illustration of the TWIST module concept for simulating tree water deficit (TWD) and relative tree water content (RWC_tree_). TWIST derives diagnostic proxies of internal water status from externally supplied inputs – transpiration (*E*), relative soil water content (*θ*_rel_), and the available tree water pool (*W*; e.g. based on Eq. 2.5). At each time step *t*, soil water uptake (*U*) compensates the fraction *F_E_*of *E* (*U_E_*) and the fraction *F*_TWD_ of the existing TWD (*U*_TWD_). When the relative soil water content (*θ*_rel_) drops below the threshold *F_θ_*, *U* is further reduced through the drought limitation function *f*_drought_. TWD decreases when *U > E* (refilling) and increases when *U < E* (depletion), resulting in changes of RWC_tree_. Blue arrows indicate water uptake from the soil, and the red arrow represents the depletion of internal water storage. Further details on variable definitions, units, and parameter values are provided in Table 1.

**Table 1:**
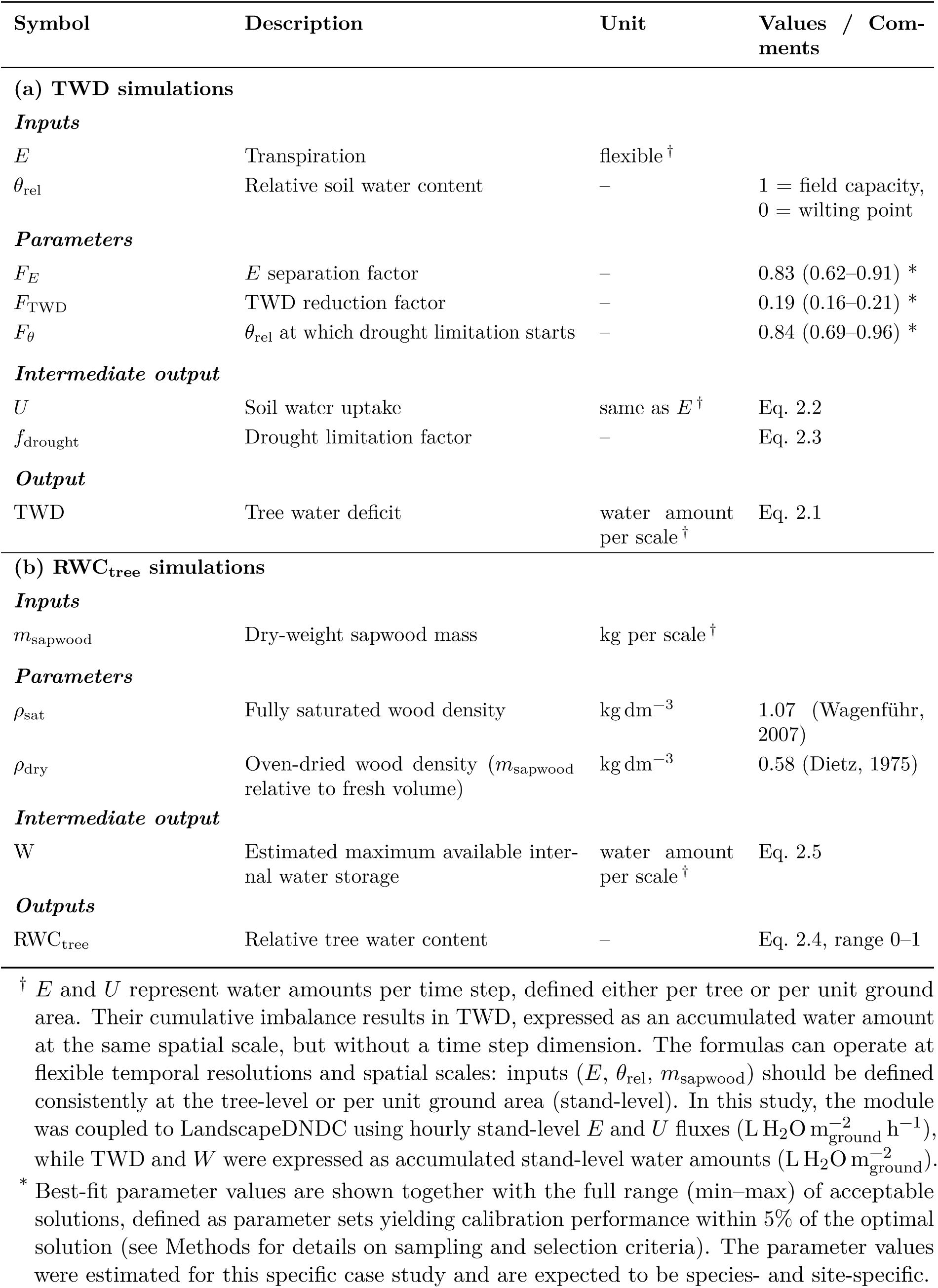
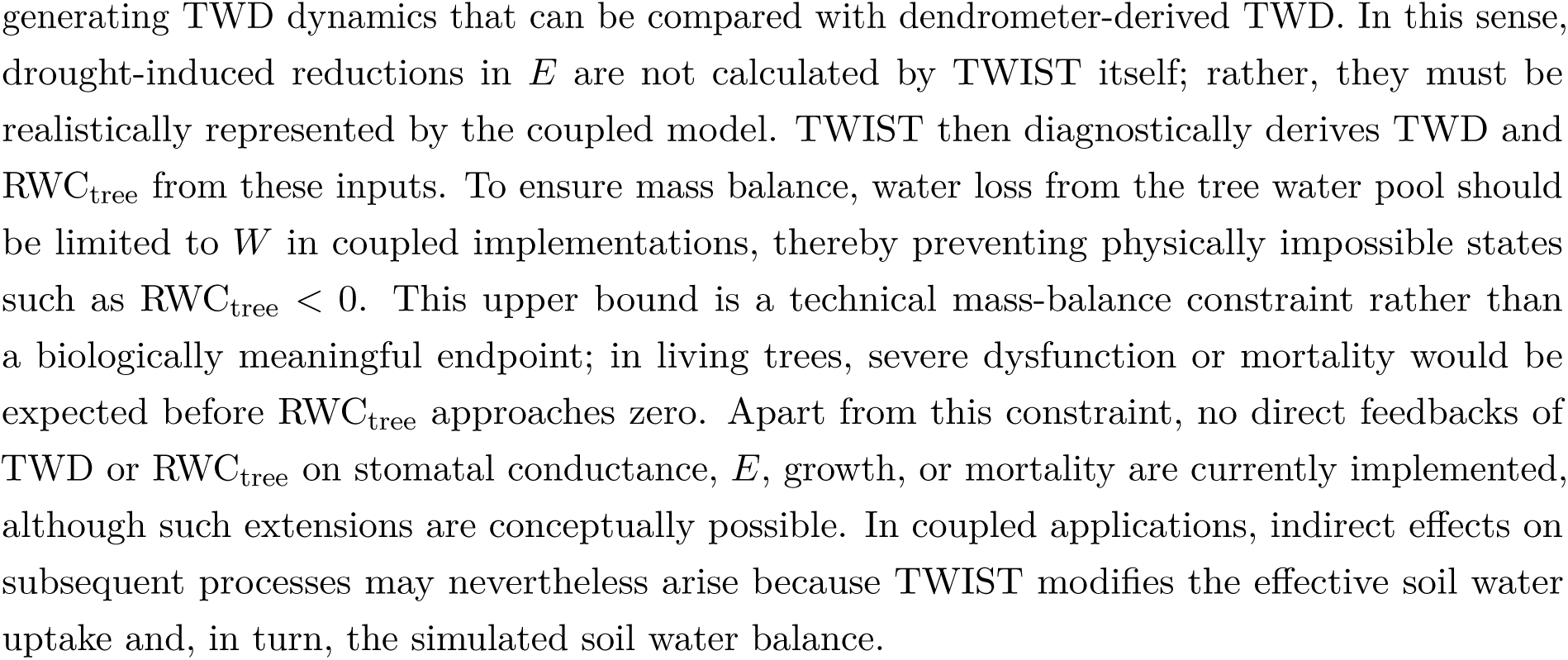
Variables and parameters used in the TWIST module to simulate (a) tree water deficit (TWD) and (b) relative tree water content (RWC_tree_) at the Štítná *Fagus sylvatica* site. Symbols, units, and brief descriptions are provided.

The module requires *E* and relative soil water content (*θ*_rel_) – ranging from 1 at field capacity to 0 at the wilting point – as inputs. Here, *E* denotes water loss through the plant pathway rather than total evapotranspiration. The formulation is flexible regarding temporal resolution, spatial scale, and units: *E* can be provided at any time step (typically sub-daily) and for individual trees or entire stands. The resulting simulated TWD represents the accumulated internal water deficit and is expressed as a volume- or mass-based water amount per tree or ground area (e.g., 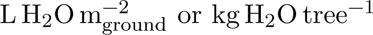). This differs from dendrometer-derived TWD, which is expressed as stem shrinkage (*µ*m) and represents a proxy for internal water deficit rather than a direct volume-based deficit.

TWD is updated at each time step (*t*) according to the previously accumulated deficit TWD*_t_*_−1_ (initialized at 0) and the current *E* and uptake rates:

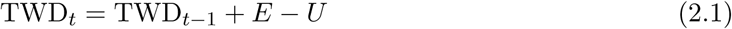

Thus, when transpiration exceeds uptake (*E > U*), TWD increases, whereas it decreases when water uptake exceeds transpiration.

The calculation of *U* forms the core of the module. It is influenced by species- and site-specific hydraulic limitations represented by three empirical parameters – *F_E_*, *F*_TWD_ and *F_θ_* – and by the drought limitation function *f*_drought_(*θ*_rel_):

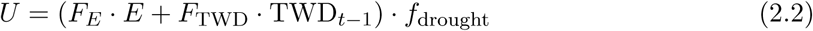

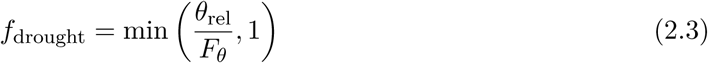

The three parameters are dimensionless and constrained between 0 and 1. *F_E_* controls the fraction of current transpiration demand that can be compensated by water uptake within the same time step. Higher *F_E_* values therefore reduce the immediate accumulation of TWD, whereas lower values increase the lag between water loss and replenishment. The parameter *F*_TWD_ controls the fraction of the previously accumulated deficit that can be replenished within a time step, thereby regulating the pace of TWD recovery. The parameter *F_θ_*defines the *θ*_rel_ below which *U* becomes additionally limited. For *θ*_rel_ *< F_θ_*, *f*_drought_ declines linearly with decreasing *θ*_rel_, resulting in direct limitation of *U*. At wilting point (*θ*_rel_ = 0), *f*_drought_ becomes zero and *U* ceases entirely. A schematic representation of all fluxes is shown in Fig. 1.

Conceptually, TWIST represents water uptake as a response to two demand components, namely *E* and TWD*_t_*_−1_, rather than resolving explicit *ψ* gradients or tissue-specific capacitances. *F_E_* and *F*_TWD_ scale how efficiently these demands can be met under non-limiting soil moisture conditions, thereby controlling the lag between water loss, uptake, and storage replenishment. They can therefore be interpreted as effective empirical controls related to resistances along the soil-root-xylem pathway and the xylem-storage pathway, but should not be mistaken for direct estimates of specific hydraulic traits. Additional limitation due to soil drying is represented through *f*_drought_(*θ*_rel_). Thus, the parameters are empirical in the present implementation, but their roles remain linked to interpretable hydraulic processes rather than acting as purely abstract fitting coefficients.

Suitable parameters for a specific site and species can be estimated with the automated calibration method described in Section 2.5. All variables, units, and parameter values used in this study are summarized in Table 1a. To examine how individual parameters influence TWD simulation behavior, a sensitivity analysis (one parameter is varied while the others are held constant) is presented in Fig. 6.

#### 2.1.2 Simulating relative tree water content

To obtain a model-derived proxy of tissue hydration, TWIST derives relative water content within the tree or stand from TWD:

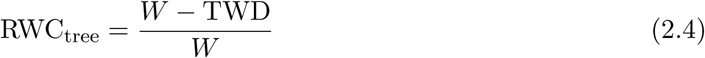

Here, *W* denotes the maximum amount of water stored in the tree or stand at full saturation. In this study, we estimate it from dry-weight sapwood mass (*m*_sapwood_) – a commonly modeled variable – and the standard density parameters *ρ*_sat_ and *ρ*_dry_, which relate water-saturated and oven-dry sapwood mass, respectively, to fresh sapwood volume:

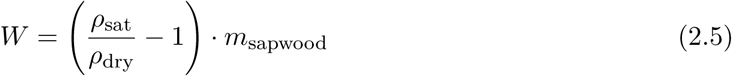

This formulation approximates the available internal water storage as the difference between saturated and dry sapwood mass (*m*_sat-sapwood_ − *m*_sapwood_), assuming that leaf, root, and core wood compartments contribute negligibly to total storage. *W* must share the same unit as TWD (1 kg H_2_O ≈ 1 L H_2_O) to ensure that RWC_tree_ remains dimensionless. All values and units used for RWC_tree_ simulations are summarized in Table 1b.

Thus, RWC_tree_ equals 1 when tissues are fully saturated and declines as TWD increases. To ensure mass balance, TWD cannot exceed the total available water storage *W*, which constrains RWC_tree_ ≥ 0. A value of 0 would correspond to complete depletion of the modeled available water storage (oven-dried conditions). RWC_tree_ therefore provides a comparable diagnostic proxy of relative tree water status over time.

#### 2.1.3 Required inputs and coupling concept

TWIST is designed as a diagnostic add-on module that can be coupled to process-based models providing *E*, *θ*_rel_, and an estimate of *W* at consistent temporal and spatial scales, thereby generating TWD dynamics that can be compared with dendrometer-derived TWD. In this sense, drought-induced reductions in *E* are not calculated by TWIST itself; rather, they must be realistically represented by the coupled model. TWIST then diagnostically derives TWD and RWC_tree_ from these inputs. To ensure mass balance, water loss from the tree water pool should be limited to *W* in coupled implementations, thereby preventing physically impossible states such as RWC_tree_ *<* 0. This upper bound is a technical mass-balance constraint rather than a biologically meaningful endpoint; in living trees, severe dysfunction or mortality would be expected before RWC_tree_ approaches zero. Apart from this constraint, no direct feedbacks of TWD or RWC_tree_ on stomatal conductance, *E*, growth, or mortality are currently implemented, although such extensions are conceptually possible. In coupled applications, indirect effects on subsequent processes may nevertheless arise because TWIST modifies the effective soil water uptake and, in turn, the simulated soil water balance.

#### 2.1.4 Idealized module behavior experiment

To illustrate the diagnostic response of TWIST under declining soil moisture and reduced *E*, an idealized simulation is presented in Fig. 2. Here, prescribed daily *E* cycles and fixed *θ*_rel_ values served as module input, while parameter values were set according to the best fit for the Štítná site. Each column represents responses under different soil water conditions, ranging from field capacity to wilting point, with gradually decreasing *E* amplitudes that resemble stomatal closure. For each time series, TWD was initialized as zero, and the module calculated *f*_drought_, *U*, and the resulting TWD for each time step according to Eqs. 2.1–2.3. This idealized module response experiment was designed to illustrate simulated TWD responses rather than to reproduce specific measurements.

**Figure 2:**
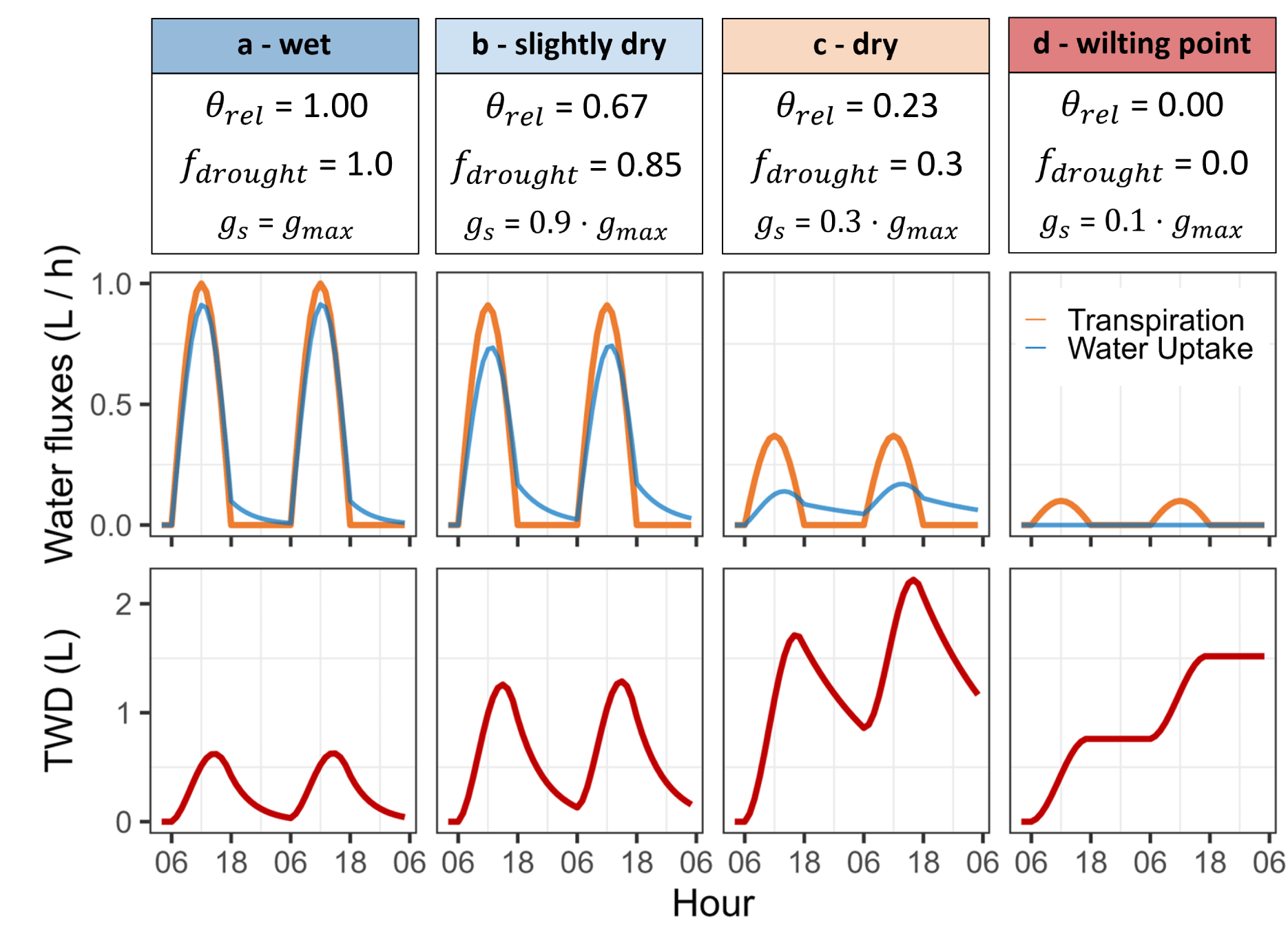
Conceptual representation of the TWIST module behavior exemplified under four increasing drought intensities (a-d). A fixed relative soil water content (*θ*_rel_), ranging from field capacity in (a) to wilting point in (d), determines the drought limitation factor (*f*_drought_; Eq. 2.3) in each column. In parallel, the amplitude of daily transpiration is reduced to resemble drought-induced reductions of stomatal conductance (*g_s_*) until its minimum *g_min_* = 0.1 · *g_max_*is reached. Vapor pressure deficit is assumed to be constant. Top panels show two diurnal courses of transpiration (orange) and the resulting soil water uptake (blue; Eq. 2.2). Imbalances between soil water uptake and transpiration result in dynamic changes in tree water deficit (TWD; red; Eq. 2.1) as shown in the bottom panels.

### 2.2 Coupling with the LandscapeDNDC ecosystem model

For this work, TWIST was integrated into LandscapeDNDC to enable multi-year stand-level simulations of TWD and RWC_tree_ for specific forest sites.

#### 2.2.1 General LandscapeDNDC properties

LandscapeDNDC is a process-based ecosystem model designed to simulate coupled carbon, water, and nitrogen fluxes between vegetation, soil, and the atmosphere at the site scale (Grote et al., 2011a; Haas et al., 2013). The model comprises a flexible, modular system and has been applied across various ecosystems, including forests, croplands, and grasslands (Grote et al., 2011b; Molina-Herrera et al., 2016; Dirnböck et al., 2020; Cade et al., 2021; Kraus et al., 2022; Sifounakis et al., 2024). It represents vegetation and soil as dynamically interacting compartments, integrating canopy microclimate, plant physiology, plant dimensional growth, and soil biogeochemistry. Simulations are initialized with actual soil- and vegetation properties (i.e., without significant spin-up periods) and are driven by meteorological forcing (temperature, vapor pressure deficit (VPD), precipitation, radiation, pressure), CO_2_ concentration, nitrogen deposition, and ecosystem management at daily to sub-daily resolution. The model calculates above- and below-ground processes within distinct canopy- and soil layers.

For each soil layer, soil water content (SWC) is simulated and normalized between the layer-specific field capacity and wilting point. A mean relative SWC (*θ*_rel_) is then derived as the weighted average across all soil layers, where weighting follows the vertical distribution of fine roots. For forest applications, Nadal-Sala et al. (2024) recently integrated tree hydraulic processes into LandscapeDNDC, enabling the simulation of soil and plant *ψ* and the calculation of transpirational water loss *E* via stomatal regulation following Eller et al. (2020). Thus, *E* is simulated dynamically in response to meteorological forcing and plant water status. In the original configuration (without the TWIST module), the resulting transpired water is removed directly from the soil layers until a species-specific soil water potential threshold is reached, at which point fine-root water uptake ceases completely (soil-to-root decoupling).

LandscapeDNDC also simulates a range of other ecosystem processes, including total evapo-transpiration, stomatal-conductance-dependent carbon assimilation, allocation of carbon and nitrogen into distinct compartments (foliage, fine roots, sapwood, and structural reserves), respiration and senescence, which decreases the living biomass of all compartments, including sapwood.

The most sensitive LandscapeDNDC parameters for photosynthesis and water balance (Table S1) were set within physiologically plausible ranges to resemble site-measured seasonal dynamics of total evapotranspiration (Fig. S1), gross primary productivity (Fig. S2), and SWC (Fig. S3).

#### 2.2.2 Integration of TWIST in LandscapeDNDC

In contrast to the original setup of Nadal-Sala et al. (2024), TWIST introduces an explicit tree water pool *W* in LandscapeDNDC (Eq. 2.5). Soil water uptake is estimated by Eq. 2.2, allowing imbalances between uptake and transpirational water loss to drive gradual depletion and refilling of this pool. The LandscapeDNDC variables *θ*_rel_ (–), 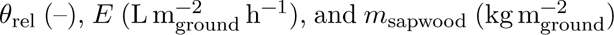 were used as inputs for the newly developed equations. Because *E* in LandscapeDNDC already responds dynamically to plant water status via stomatal regulation, TWIST is driven by a drought-responsive rather than unconstrained *E* signal. To prevent physically impossible states, depletion of the tree water pool was limited to the available storage *W*, such that TWD cannot exceed *W* and RWC_tree_ cannot fall below zero.

### 2.3 Evaluation site

TWIST was evaluated during seven consecutive years (2018–2024) at the Štítná forest site, located in the Czech Republic (49^◦^02^′^09.510^′′^ *N,* 17^◦^58^′^11.640^′′^ *E*; 550 *m* above sea level). The stand consists of European beech trees (*Fagus sylvatica L.*; 286 trees ha^−1^; DBH = 37 ± 13 *cm*; height = 31 ± 7 *m* (mean ± std; measured 2018)). At the site, meteorological measurements, eddy covariance ecosystem flux measurements, SWC measurements, and stem diameter measurements were conducted. All meteorological and ecosystem exchange measurements were conducted above the forest canopy at 44 *m* height. The eddy covariance system consists of a Gill ultrasonic anemometer (HS-50, Gill Instruments, UK) and a LI-COR infrared gas analyzer (LI-7200, LI-COR, NE, USA). Processing of raw 20 Hz data was performed by EddyPro Software (LI-COR, NE, USA) with settings described in McGloin et al. (2019). Quality-controlled half-hourly products were post-processed using the standardized ONEFlux processing pipeline (Pastorello et al., 2020) by the FLUXNET network.

Half-hourly SWC measurements were performed at 5, 10, and 30 cm soil depths using ThetaProbe sensors (ML2x 355, Delta-T, UK).

Automatic, high-resolution band dendrometers (DRS26; EMS Brno, Czech Republic; 1 *µm* resolution) were used to monitor changes in stem circumference of five individuals (DBH = 46 ± 10 *cm*; height = 33 ± 3 *m* (mean ± std)). The sensors were installed approximately 1.3 m above ground level. Stem dynamics were recorded continuously at half-hourly intervals on trees in the co-dominant and dominant canopy layers within the eddy-covariance tower footprint. Circumference measurements were converted to changes in stem radius. To isolate water-related, reversible stem variations, we applied the zero-growth concept (Zweifel et al., 2016), which assumes that irreversible radial growth occurs only when new stem-radius maxima are reached, while short-term deviations below these maxima reflect reversible water-related shrinkage. This approach enabled direct calculation of observed TWD (stem shrinkage; *µ*m).

To avoid frost-related artifacts in the observed TWD data (Zweifel and Häsler, 2000), an individual growing-season period was defined for each year. The start of the season was set to the first day after 1 April on which TWD returned to 0 (indicating full recovery from winter effects), and the end of the season to the last day before the daily mean air temperature dropped below 5^◦^C.

### 2.4 Normalization of TWD

To compare the temporal dynamics of observed and simulated TWD (TWD_obs_ and TWD_sim_, respectively), all values were normalized following the procedure of Peters et al. (2025). For the dendrometer measurements, maximum daily shrinkage (MDS) was calculated for each tree and each day as the difference between the daytime maximum TWD_obs_ (10 am–11 pm) and the pre-dawn minimum TWD_obs_ (4–8 am), and the 99th percentile of all daily MDS (MDS*_q_*_99_) was determined. Next, the TWD_obs_ time series of each tree was divided by its MDS*_q_*_99_, yielding a normalized TWD (TWD_norm_) per individual. Across the seven measurement years and across all five trees, the average MDS*_q_*_99_ was 33.2 ± 2.3 *µm* (mean ± sd). The mean and standard deviation of all individual TWD_norm_ trajectories were used for further analysis.

For TWD_sim_, an equivalent normalization was applied using the 99th percentile of the maximum daily water loss (MDWL*_q_*_99_; volume-based analogue of MDS) across the study period, resulting in MDWL*_q_*_99_ = 1.01 L m^−2^ (equivalent to 35.0 L tree^−1^).

According to Peters et al. (2025), this normalization enables the identification of relative drought stress thresholds and facilitates comparison of tree water status across individuals and species, independent of absolute stem size or shrinkage magnitude. Applied to TWD_sim_, it also bridges differences in units (water-volume deficit vs. stem shrinkage) and spatial scales (stand-level vs. tree-level), thereby enabling comparison of tree-level dendrometer signals with stand-level model dynamics.

### 2.5 Parameter calibration and model evaluation

The three TWIST parameters were calibrated against the temporal dynamics of Štítná dendrometer-derived TWD observations in 2018, while 2019–2024 were used for independent evaluation. 2018 was selected as a calibration year because its second half was exceptionally warm and dry at the Štítná site (Nezval et al., 2025), while its first half also included extended periods without strong SWC limitation (Fig. S3), overall providing a wide range of conditions. Parameter estimation was restricted to bounded intervals of 0–1 for *F_E_* and *F*_TWD_, and 0.01–1 for *F_θ_*. For each tested parameter set, TWIST was run over the full simulation period (2018–2024). Only time steps with available observations were used for scoring. Simulated and observed TWD were each normalized over the full available time series, as described above. The calibration objective combined the root mean square error (RMSE) between simulated and observed TWD_norm_ at hourly resolution with the RMSE of daily mean TWD_norm_, thereby constraining both sub-daily dynamics and day-to-day variation. To give both components comparable influence, hourly and daily RMSE were scaled by the corresponding errors of a fixed reference parameter set (*F_E_* = 0.6, *F*_TWD_ = 0.3, *F_θ_* = 0.7), used solely as an internal scaling baseline, and weighted equally. The optimization was performed with a bounded L-BFGS-B algorithm, a method that efficiently handles box-constrained parameter spaces (Byrd et al., 1995), using a multi-start approach with 10 initial parameter sets, including the reference parameter set and 9 randomly sampled starts within the parameter bounds. The best-fit solution was defined as the parameter set yielding the minimum combined objective value during the calibration period.

Model evaluation was performed independently for the 2019–2024 growing seasons using the calibrated best-fit parameter set without further adjustment. Performance was quantified at both hourly and daily resolution using RMSE, bias, and *R*^2^ between simulated and observed TWD_norm_. To further characterize parameter robustness and equifinality, the bounded parameter space was additionally explored by uniform random sampling of 8000 parameter combinations. Each sampled set was evaluated with the same objective function as used for calibration. Acceptable solutions were defined as all parameter sets with an objective value within 5% of the optimized best fit. The corresponding parameter distributions are reported in Table S2, and all acceptable parameter sets are listed in Table S3. For reproducibility and new applications, the calibration R code is provided (see Data and code availability).

### 2.6 Rainfall-suppression stress test

To explore TWIST behavior under imposed soil water depletion, an idealized stress test was simulated by suppressing rainfall after the 2018 TWD peak in mid-August at the Štítná site. All other meteorological drivers (temperature, radiation, and VPD) were prescribed as a repetition of the observed weather conditions during the peak drought period (14–23 August; Fig. 7).

This setup produced continuous soil moisture depletion, allowing us to examine how TWD and RWC_tree_ evolve when residual water loss persists in the absence of soil water recharge. The experiment was therefore used as a technical boundary-case test of model behavior, not as a realistic drought scenario.

All data processing, statistical analyses, and visualizations were conducted in R (R Core Team, 2022).

## 3 Results

### 3.1 Theoretical TWIST module behavior

The theoretical example presented in Fig. 2 illustrates how TWIST represents the transition from complete to incomplete nocturnal TWD recovery across the soil moisture gradient. Under well-watered conditions (Fig. 2a), the simulated uptake did not fully cover *E* during the day, resulting in an increase in TWD, but the deficit could be fully replenished during the night. As *θ*_rel_ declined and drought limitation began to constrain uptake (*f*_drought_ *<* 1, Fig. 2b-d), an increasing fraction of *E* could no longer be balanced by *U*, causing higher daytime TWD peaks and slower replenishment.

At moderate soil dryness (Fig. 2b), TWD still returned towards zero at night, but recovery took several hours longer compared with wet conditions. When *θ*_rel_ dropped further (Fig. 2c), storage refilling was incomplete and a residual TWD persisted into the following day. This carry-over deficit marks the onset of TWD accumulation, where daily minimum TWD progressively increases despite reduced *E*. Under extreme drought (Fig. 2d), soil water uptake ceased entirely (*U* = 0), and the minor residual *E* occurring despite stomatal closure directly translated into continuous TWD increase.

### 3.2 TWIST module evaluation

To assess model performance, TWIST was coupled to the ecosystem model LandscapeDNDC and applied to the Štítná beech forest site (Fig. 3a). Parameters were optimized based on the 2018 growing season and evaluated across the remaining years. For the calibration year, the best-fit parameter set was derived by minimizing the mismatch between simulated and observed TWD (Section 2.5). The resulting simulation shows close agreement in both temporal dynamics and relative amplitude (Fig. 3b). The timing of drought onset, peak, and recovery was well captured, resulting in high agreement (hourly *R*^2^ = 0.92) and low error (hourly RMSE = 0.24). Minor deviations in normalized peak amplitudes and the magnitude of diurnal cycles occurred, but the low overall hourly bias of −0.06 indicates no strong systematic over- or underestimation during the calibration period.

**Figure 3:**
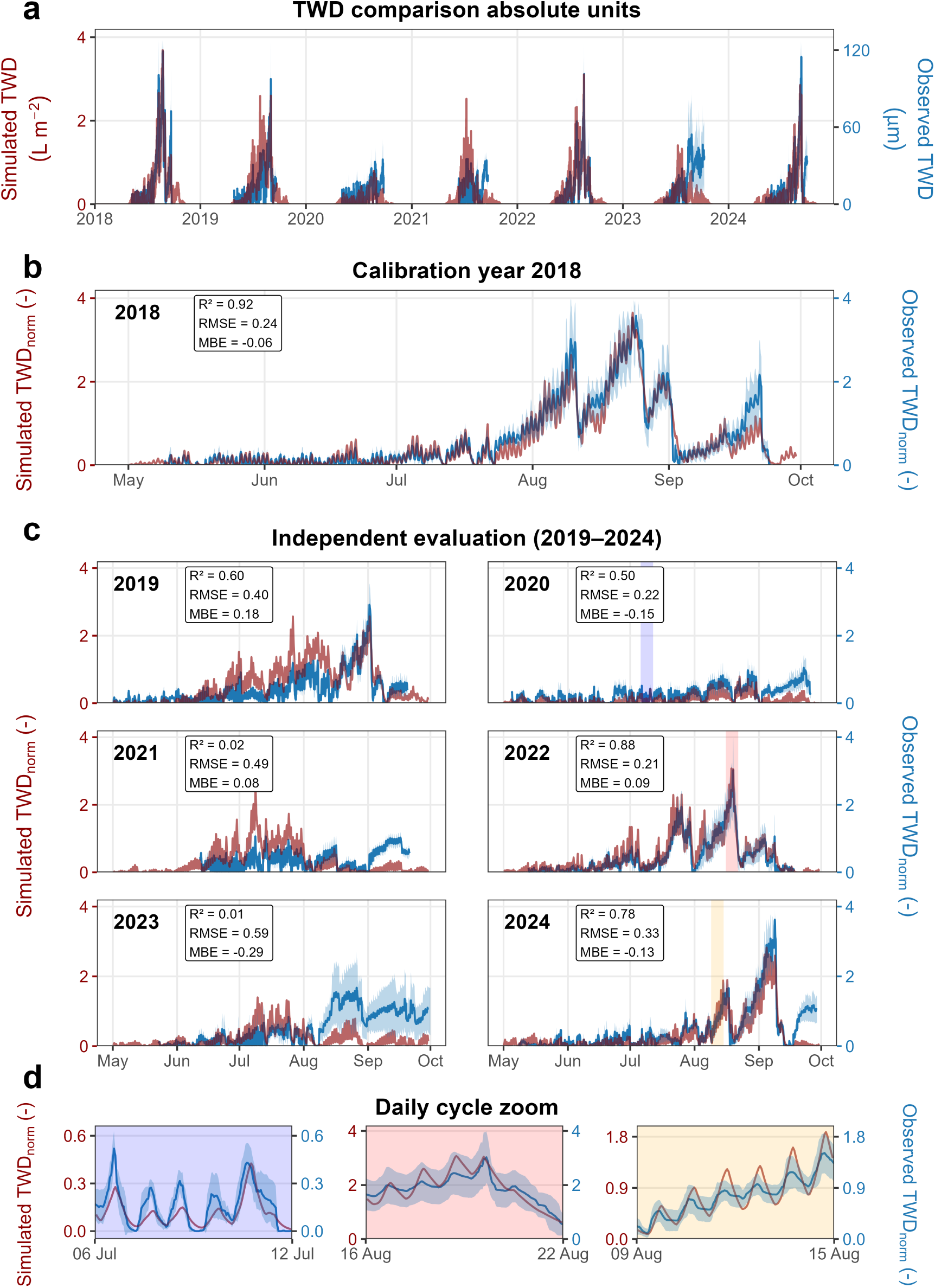
Evaluation of simulated tree water deficit (TWD; red) against dendrometer-based TWD observations (blue; *n* = 5, mean ± sd) at the Štítná forest site for the years 2018–2024. (a) shows absolute values for context, where simulated TWD represents the derived water volume deficit at the stand-level (L m^−2^ ground area), while observed TWD denotes stem diameter shrinkage at the tree-level (*µ*m). (b) shows the calibration year 2018, for which the three TWIST parameters were optimized by minimizing the RMSE of hourly values and daily means between simulated and observed normalized TWD (hourly shown; daily values see Table 2). (c) presents independent evaluation for the years 2019–2024 using the calibrated best-fit parameter set. (d) zooms into the color-coded periods highlighted in (c), illustrating short-term TWD dynamics.

**Table 2:** Model performance metrics for calibration (2018) and independent evaluation (2019—2024) periods, based on hourly and daily aggregated data. RMSE denotes the root mean square error, calculated from either hourly values or daily means of normalized tree water deficit, Bias denotes the mean error, and *R*^2^ the coefficient of determination.

| Metric | Calibration (2018) | Evaluation (2019–2024) |
| --- | --- | --- |
| Hourly RMSE (–) | 0.24 | 0.40 |
| Daily RMSE (–) | 0.20 | 0.39 |
| Hourly Bias (–) | -0.06 | -0.05 |
| Daily Bias (–) | -0.06 | -0.05 |
| Hourly $R^2$ (–) | 0.92 | 0.42 |
| Daily $R^2$ (–) | 0.94 | 0.42 |

This calibrated parameter set was subsequently applied without further adjustment to the independent evaluation period 2019–2024 (Fig. 3c). During this evaluation period, the model captured the general normalized TWD dynamics across contrasting conditions, although performance varied between years. Good agreement was observed for years such as 2019, 2022, and 2024, whereas lower agreement was observed in 2021 and 2023. This variability is reflected in reduced overall performance metrics compared to the calibration period (e.g., hourly *R*^2^ = 0.42, RMSE = 0.40, bias = −0.05; Table 2).

At the sub-daily scale, the model reproduced the characteristic diurnal dynamics of water depletion during daytime and rehydration during nighttime (Fig. 3d). The amplitude of these fluctuations was not always reproduced exactly, but deviations did not show a consistent directional bias across years. Instead, both over- and underestimation occurred depending on period and year. Nevertheless, the overall temporal structure was captured consistently across both wet and dry periods. In particular, the transition from complete nocturnal recovery under moist conditions to persistent deficits under drought was well represented. Together with the comparison of hourly and daily performance metrics (Table 2), this indicates that the model captures both short-term fluctuations and the broader day-to-day development of TWD reasonably well.

Overall, the results indicate that the parameter set calibrated for 2018 captured the main TWD patterns across subsequent years, albeit with lower performance than in the calibration year.

### 3.3 TWD and RWC_tree_ dynamics at the site

The calibrated parameter set was used to analyze the temporal dynamics of TWD and RWC_tree_ at the site (Fig. 4). Periods of high *E* combined with declining *θ*_rel_ led to a progressive accumulation of TWD, while rainfall events and increasing *θ*_rel_ triggered recovery of internal water reserves. This resulted in a pronounced seasonal cycle, with TWD peaks during dry summer periods indicating increased internal water depletion. The highest peak occurred in 2018 (∼ 3.6 L m^−2^). Similarly high peaks were simulated in 2019, 2021, 2022, and 2024 (∼ 2.6–3.1 L m^−2^), whereas substantially lower values were observed in 2020 (∼0.8 L m^−2^) and 2023 (∼1.4 L m^−2^).

**Figure 4:**
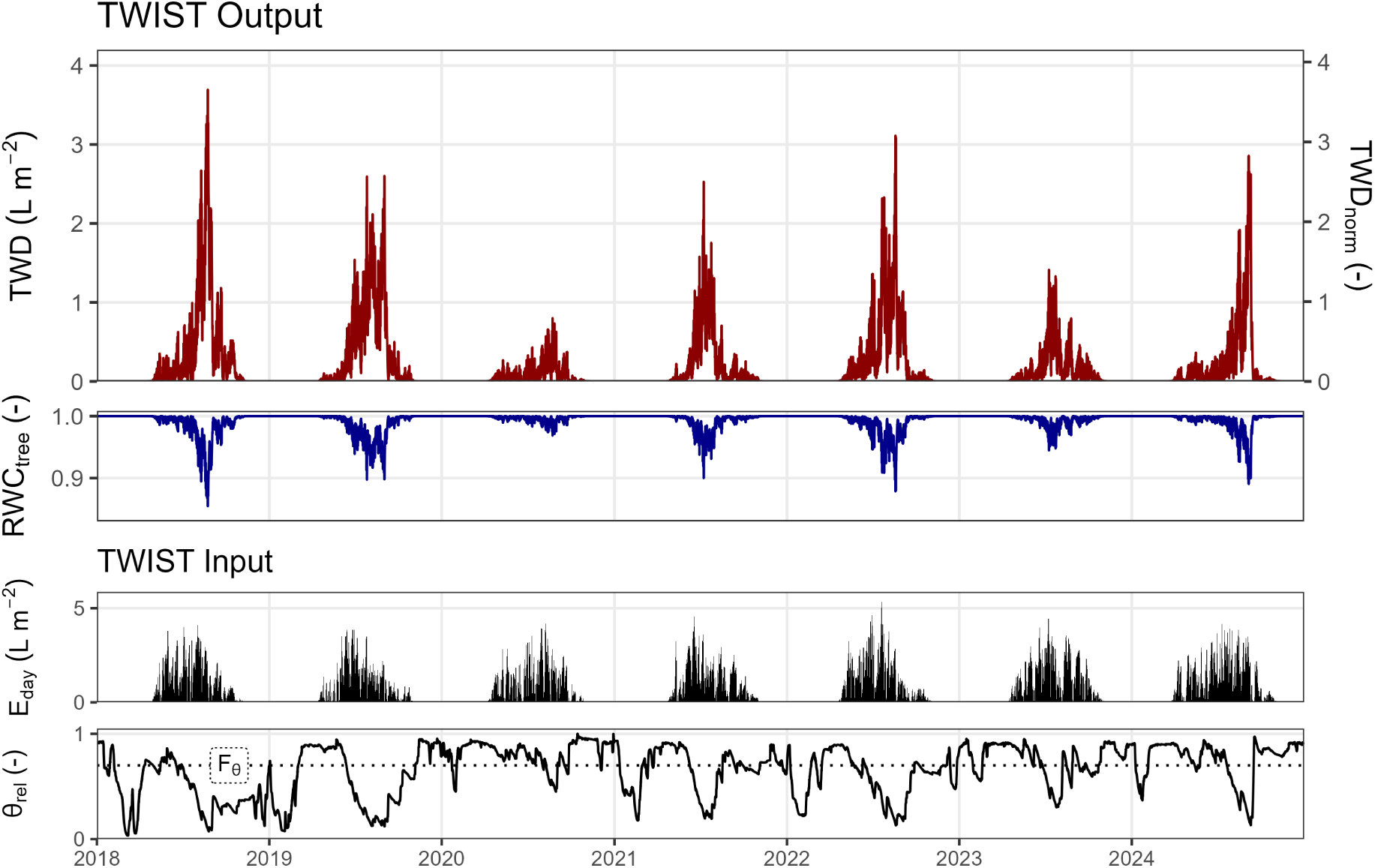
Seasonal dynamics of simulated tree water deficit (TWD) and relative tree water content (RWC_tree_) at the Štítná forest site 2018–2024. Hourly TWD is shown both as absolute values (left axis; liters per *m*^2^ ground) and as normalized deficit (right axis; for normalization procedure see Section 2.4). Hourly transpiration (aggregated to daily values, *E*_day_) and relative soil water content (*θ*_rel_; 1 = field capacity, 0 = wilting point) were simulated by the LandscapeDNDC model and used as inputs for the TWIST module. In the *θ*_rel_ panel, the *F_θ_* threshold indicating the onset of additional soil water uptake limitation is shown as a dotted line.

The complementary RWC_tree_ signal closely mirrored TWD, expressing the relative depletion of internal water storage (on average ∼25.3 L m^−2^; equivalent to ∼884 L tree^−1^). RWC_tree_ remained close to 1 under moist conditions (e.g., 2020), and declined to approximately 0.86 during the drought year 2018, reflecting partial tissue dehydration. Together, TWD and RWC_tree_provided a consistent representation of tree water dynamics and its seasonal variability at the site.

### 3.4 Parameter space and equifinality

Building on the calibrated parameter set used in the model evaluation, the parameter space was further explored to assess parameter uncertainty and equifinality. This exploration showed that only a small subset of parameter combinations yielded performance comparable to the best-fit solution (Fig. 5a). Acceptable performance occurred only within a narrow region of the sampled parameter space, indicating that good model performance was restricted to specific parameter combinations rather than broadly distributed across the parameter domain. The parameters *F_E_*and *F_θ_* exhibited comparable ranges, with acceptable values of *F_E_* = 0.83 (0.62 – 0.91) and *F_θ_* = 0.84 (0.69 – 0.96), respectively (best-fit value and full min–max range of acceptable solutions; see also Table S2). In contrast, *F*_TWD_ was more tightly constrained, with acceptable values of 0.19 (0.16 – 0.21).

**Figure 5:**
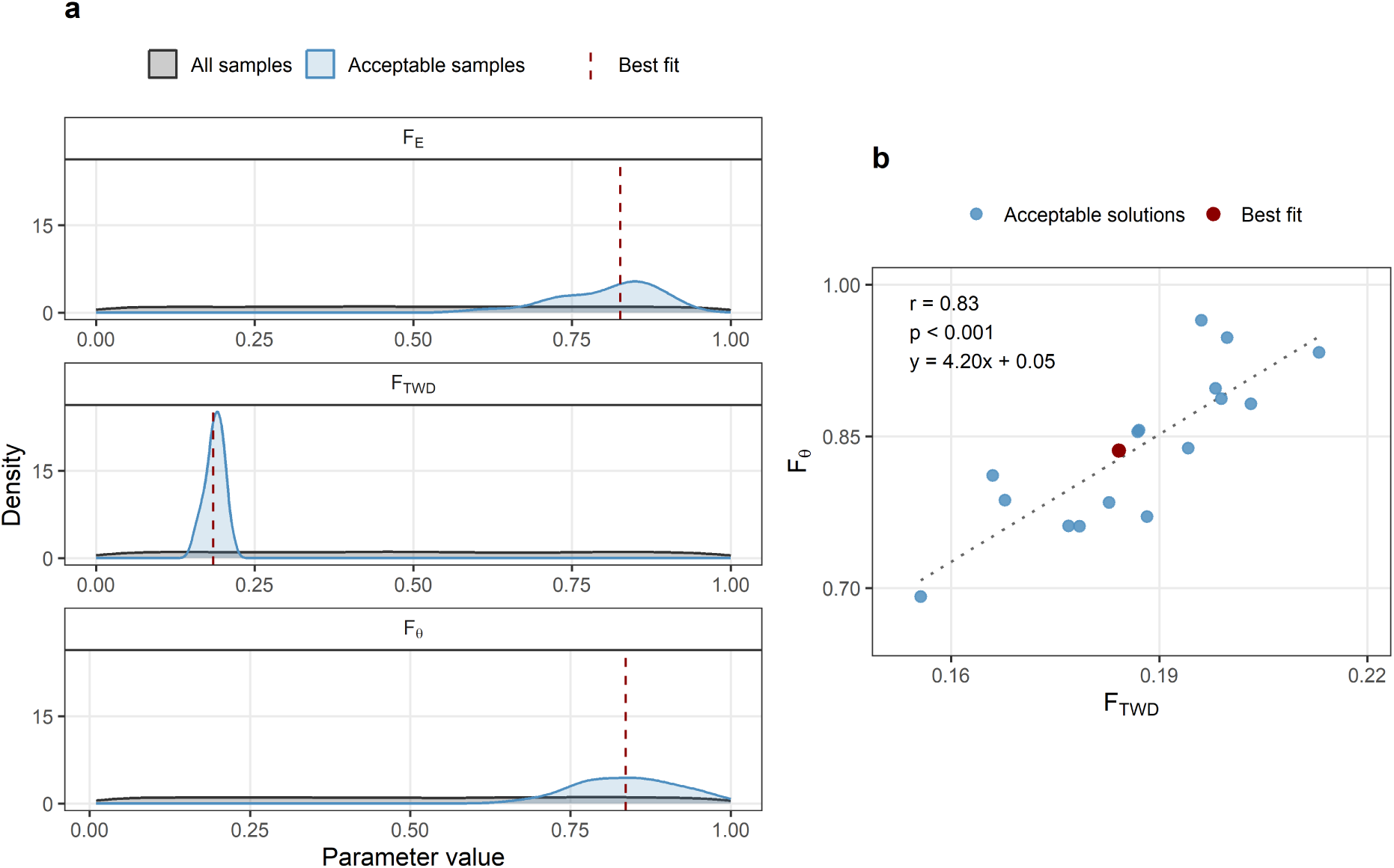
Parameter calibration and uncertainty analysis of the three TWIST model parameters based on the 2018 calibration period. (a) Distributions of all sampled parameter combinations (uniform sampling within the parameter bounds; *n* = 8000; grey) and the subset of acceptable solutions (*n* = 17; blue), defined as parameter sets with an objective value within 5% of the best-fit objective value. Dashed vertical lines indicate the best-fit parameter values. (b) Relationship between *F*_TWD_ and *F_θ_*. Blue points represent acceptable parameter combinations, and the red point indicates the best-fit solution. The dashed line shows a linear regression fitted to the acceptable solutions. The annotated value denotes the Pearson correlation coefficient (*r*) between *F*_TWD_ and *F_θ_* across all acceptable solutions, indicating the strength and direction of their linear relationship. Similar relationships including the *F_E_* parameter are shown in Fig. S4.

Within this constrained parameter space, a clear relationship emerged between *F*_TWD_ and *F_θ_* (Fig. 5b). Acceptable solutions, i.e. parameter sets yielding model performance within 5% of the best-fit solution, followed a distinct positive linear trend (*r* = 0.83, *p <* 0.001), indicating that combinations of higher *F*_TWD_ were associated with higher *F_θ_* values. A weaker but significant positive relationship was also found between *F_E_* and *F*_TWD_ (Fig. S4; *r* = 0.50, *p* = 0.039), suggesting partial parameter compensation within the set of acceptable solutions. In contrast, no significant relationship was detected between *F_E_* and *F_θ_* (Fig. S4).

These patterns illustrate that acceptable model performance is confined to a limited and structured region of the parameter space.

### 3.5 Parameter sensitivity

The sensitivity analysis highlighted distinct roles of the three TWIST parameters in controlling TWD_sim_ dynamics under contrasting moisture conditions (Fig. 6).

**Figure 6:**
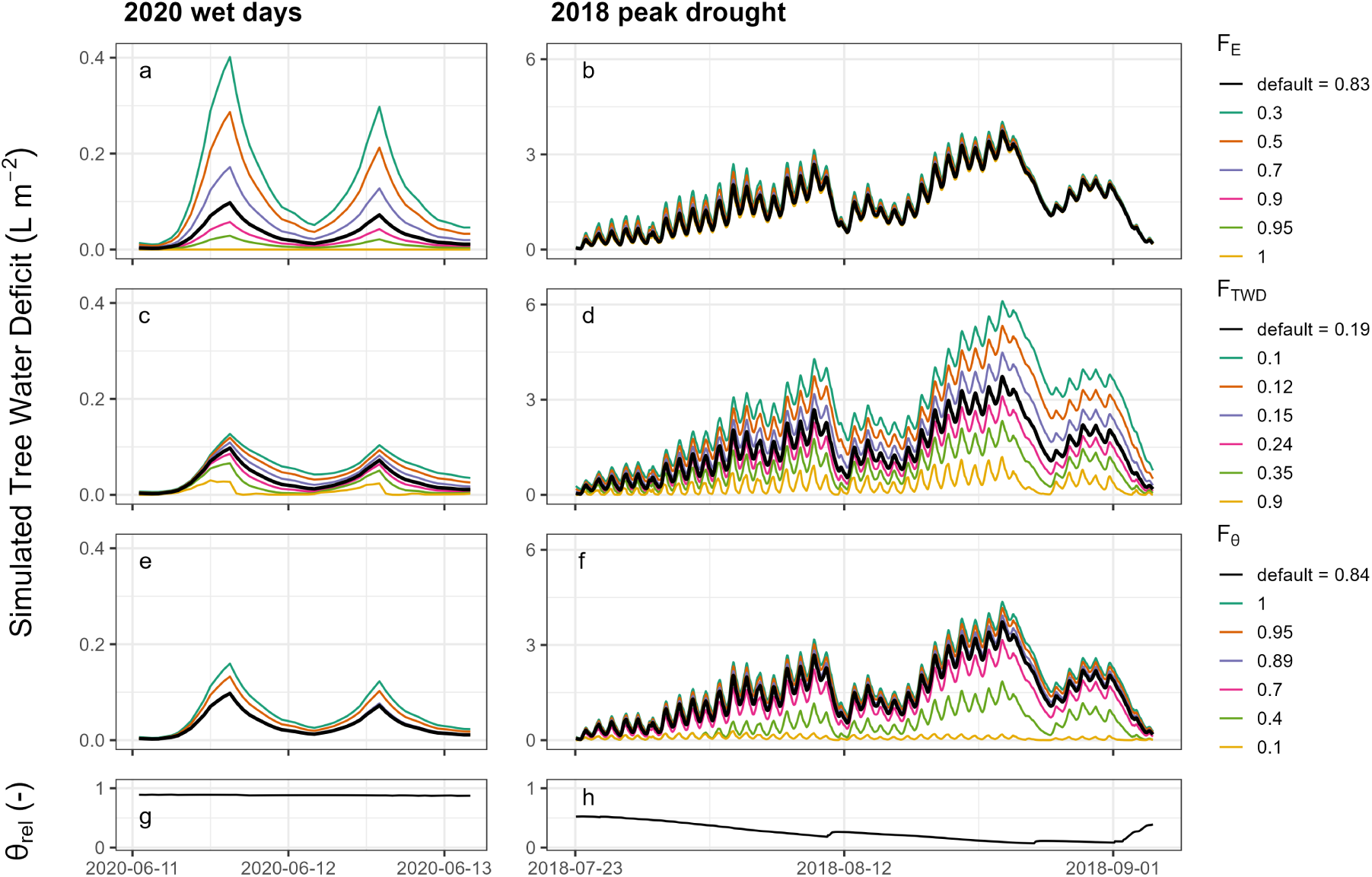
Sensitivity analysis of the three TWIST parameters *F_E_* (a,b), *F*_TWD_ (c,d), and *F_θ_* (e,f) for two exemplary wet days in 2020 (a,c,e,g) and the 2018 peak drought period (b,d,f,h) at the Štítná beech forest site. Each parameter was varied individually while the remaining parameters were kept at their site-specific best-fit values (*F_E_* = 0.83, *F*_TWD_ = 0.19, *F_θ_* = 0.84). The model inputs consist of transpiration and relative soil water content (*θ*_rel_; g,h) simulated by LandscapeDNDC.

The parameter *F_E_*primarily controlled the amplitude of daily TWD cycles. Lower values led to stronger daytime depletion and higher diurnal peaks, while higher values reduced the magnitude of these fluctuations. In contrast, *F_θ_* governed the response under prolonged dry conditions by regulating the onset and strength of soil moisture limitation. Lower *F_θ_* values delayed the reduction of water uptake and thus maintained lower TWD levels, whereas higher values enhanced drought limitation and promoted stronger deficit accumulation. The parameter *F*_TWD_ influenced both short- and long-term dynamics. Lower values reduced the efficiency of TWD compensation, resulting in slower recovery, higher daily peak amplitudes, and greater deficit accumulation under drought. Higher values accelerated rehydration and dampened both diurnal variability and longer-term TWD build-up.

### 3.6 Rainfall-suppression stress test

To examine TWIST behavior under imposed water depletion, a rainfall-suppression stress test was simulated (Fig. 7). This experiment was not intended to represent a fully realistic drought trajectory, especially after RWC_tree_ fell below 0.6, where mortality risk would be expected to increase (Sapes and Sala, 2021; Trifilò et al., 2023). Instead, it was used to test how TWIST outputs evolve under boundary-case conditions. In contrast to the observed 2018 conditions, where rainfall in late August partially restored *θ*_rel_, soil moisture in the test simulation declined continuously and asymptotically approached the wilting point (*θ*_rel_ = 0). During the first days of the scenario period, minor nocturnal TWD replenishment still occurred due to water uptake from the soil, further reducing *θ*_rel_. With increasing dryness, soil water uptake became progressively restricted, resulting in a steady increase in TWD driven by residual *E*. This behavior corresponds to phase (d) of the theoretical response illustrated in Fig. 2.

**Figure 7:**
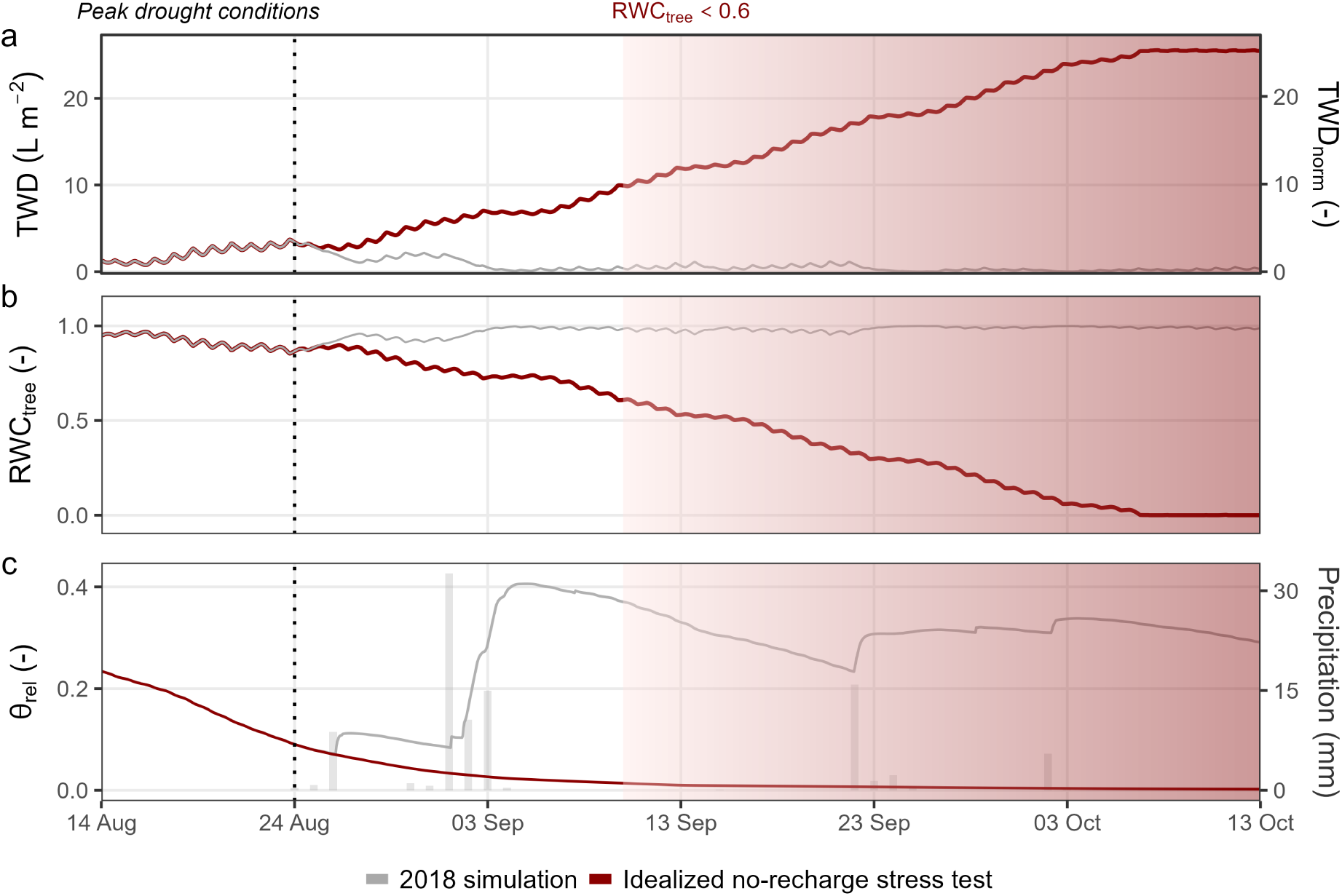
Simulated tree and soil water dynamics at the Štítná site during an idealized rainfall-suppression stress test. The baseline simulation (grey) is driven by the observed 2018 meteorological forcing, while the test simulation (dark red) artificially extends the dry-period forcing by looping the last observed dry-period weather conditions (14–23 August; peak drought conditions). The vertical dashed line marks the start of the imposed test forcing, defined by the timing of the highest TWD peak observed in 2018 (24 August). Panels show (top to bottom): (a) tree water deficit (TWD; absolute values on the left axis, normalized values on the right axis), (b) relative tree water content (RWC_tree_), and (c) relative soil water content (*θ*_rel_; left axis) with daily precipitation (bars; right axis). The shaded red area indicates the period during which RWC_tree_ falls below 0.6, shown as an indicative reference threshold associated with severe plant water stress in previous studies; dynamics beyond this range are shown to illustrate numerical model behavior rather than realistic dynamics of living trees.

RWC_tree_ mirrored the TWD trajectory, showing a continuous decline throughout the simulation. By September 9, RWC_tree_ approached 0.6, and by 7 October, it approached 0, representing the lower technical bound for internal water storage. Once this lower bound was reached, further depletion was no longer allowed, and TWD_sim_ therefore reached a plateau.

## 4 Discussion

This study shows that dendrometer-observable TWD dynamics can be represented in process-based models using a simple water-balance framework. TWIST does not replace detailed hydraulic models, but provides a model-compatible diagnostic state variable that can be compared with dendrometer-derived TWD dynamics.

### 4.1 Process-based simulation of TWD and its physiological meaning

TWD reflects the transient imbalance between *E* and water uptake, thereby encoding key information on internal tree water status and drought progression. Dendrometer-derived and simulated TWD are closely linked but differ in their physical representation: dendrometer signals quantify stem shrinkage as an observation-based proxy for water depletion in elastic tissues (Steppe et al., 2006), whereas TWD_sim_ is a model state variable representing actual volumetric water deficit across tree tissues or at the stand-scale. Although these quantities are not directly equivalent, they are expected to track the same underlying depletion–rehydration dynamics and are therefore comparable in their temporal patterns.

A more mechanistic representation of plant hydraulics could, in principle, simulate TWD and RWC_tree_ in greater detail and resolve transport pathways and storage processes across compartments, as done, for instance, in models such as SurEau and FETCH4 (Cochard et al., 2021; Ruffault et al., 2022; Missik et al., 2025). Likewise, hydraulically enhanced ecosystem models such as ED2-Hydro, ORCHIDEE-CAN-HYD and FATES-HYDRO explicitly represent tissue water storage and hydraulic traits within broader vegetation dynamics (Xu et al., 2016; Yao et al., 2022; Xu et al., 2023). These approaches offer important physiological detail, but also require substantially more parameters and a more detailed representation of hydraulic compartments and traits, making them increasingly difficult to constrain and transfer across sites and species. TWIST therefore occupies an intermediate position: it is less mechanistic than full hydraulic models, but easier to constrain and directly comparable with widely available dendrometer observations.

We show that TWIST can represent different stages of drought severity (Fig. 2). The module reproduces the characteristic diurnal pattern of daytime TWD increase followed by nocturnal recovery (De Schepper et al., 2012). As soil moisture declines, daily TWD amplitudes increase (Fig. 2b), indicating emerging soil water limitation (Peters et al., 2025). Progressive drying prolongs the replenishment of nocturnal storage, reducing the time available for stem growth (Zweifel et al., 2021b). With further drying, incomplete night-time recovery (Fig. 2c) marks the transition to full growth cessation. Residual pre-dawn TWD is then closely linked to sharp declines in midday stomatal conductance (Ziegler et al., 2024; Peters et al., 2025), underscoring its value as a robust drought indicator. As stomatal closure intensifies, *E* decreases, and daily TWD amplitudes decline (Peters et al., 2025). Under severe drought (Fig. 2d), soil water uptake approaches zero, effectively disconnecting roots from the surrounding soil matrix (Nadal-Sala et al., 2024), while continued dehydration from residual water loss (Duursma et al., 2019) further increases TWD. By reproducing these characteristic TWD patterns, TWIST provides a diagnostic representation of physiologically meaningful water-deficit dynamics that are typically absent from ecosystem models (Martinez-Vilalta et al., 2019).

### 4.2 Evaluation of TWD and RWC_tree_ simulations and associated uncertainties

Applied to field conditions, TWIST captured the main temporal patterns of TWD across the calibration and evaluation periods (Figs. 3, 4). Agreement was highest in 2018, as expected, because this year was used for calibration. During 2019–2024, performance declined but the main dynamics were still reproduced without further parameter adjustment. Notably, years with pronounced TWD peaks were reproduced more consistently than wetter years with only weak depletion signals, suggesting that stronger drought responses are more clearly constrained by the coupled setup.

A major source of uncertainty arises from the driving inputs. Errors in estimating *E* or *θ*_rel_ directly affect the simulated timing and magnitude of TWD, including both seasonal peak values and sub-daily dynamics. In the present setup, both TWIST input variables were simulated by LandscapeDNDC. The underlying LandscapeDNDC water balance was evaluated against observations of seasonal total evapotranspiration (Fig. S1) and SWC (Fig. S3). The model matched these observations reasonably well, although deviations remained. Uncertainty in *θ*_rel_ is likely particularly important, because this variable must condense vertically heterogeneous soil water availability into a single quantity that reflects the effective accessibility of soil water to the stand. Errors in *E*, in turn, propagate directly into the amplitude of diurnal TWD cycles, and a combination of low *θ*_rel_ and high *E* can contribute to exaggerated TWD accumulation. Since TWD_sim_ integrates these drivers, uncertainties in input forcing and TWIST parameterization cannot be fully disentangled here and should instead be regarded as part of the performance of the LandscapeDNDC-TWIST system as a whole. In principle, TWIST could also be driven more directly by measurements. In practice, however, deriving the required inputs consistently is not straightforward, because *E* must be separated from ecosystem evapotranspiration and SWC must be converted into *θ*_rel_ referenced to field capacity and wilting point. Process-based ecosystem models are therefore useful because they provide these quantities in a temporally and structurally consistent form. Additional measurements, particularly sap-flux-based *E* estimates and more detailed soil moisture observations, could further constrain these uncertainties.

The calibrated parameter space was constrained but not unique, indicating that the three TWIST parameters are identifiable within a limited range while still allowing partial equifinality (Fig. 5). The sensitivity analysis further revealed distinct functional roles of the parameters (Fig. 6). *F_θ_* and *F*_TWD_ mainly governed the seasonal evolution of drought stress, as reflected in their relatively high rank correlations with model performance (*ρ* = −0.61 and 0.29, respectively; Table S2). The strong coupling between these two parameters indicates that they are not independently constrained, but jointly determine model behavior. In contrast, *F_E_*primarily controlled the amplitude of daily TWD cycles and showed only a weak relationship with overall model performance (*ρ* = −0.05). This is consistent with the observed model behavior, where sub-daily TWD amplitudes were sometimes over- or underestimated (Fig. 3d), making *F_E_*comparatively challenging to constrain. The significant relationship between *F_E_* and *F*_TWD_ (Fig. S4) indicates partial interdependence, as *F*_TWD_ also influences diurnal amplitudes. Overall, these results show that suitable combinations of *F_θ_* and *F*_TWD_ reproduce the seasonality of TWD well, while *F_E_* primarily influences sub-daily TWD dynamics.

Although the parameters used in TWIST are empirical and not directly measurable, they can be related to physiological and anatomical properties of the soil–plant water transport system. In conceptual terms, *F_E_*controls the lag between current transpirational water loss and compensating soil water uptake, and may therefore be linked to vertical resistance along the xylem pathway. *F*_TWD_ controls the efficiency with which previously accumulated deficits are replenished, and may relate to radial and axial resistances between conductive xylem and storage compartments. *F_θ_* determines the onset of additional soil moisture limitation and is likely influenced by rooting depth, soil properties, and the effective accessibility of soil water. These interpretations should not be understood as direct estimates of specific hydraulic traits; however, they indicate that the calibrated parameters are linked to physiologically interpretable roles.

Two structural assumptions are particularly important and associated with further uncertainty: the shape of the drought limitation function *f*_drought_ and the estimate of the available internal water pool *W*. First, the choice of *f*_drought_ (Eq. 2.3) remains simplified, and alternative formulations may be worth testing. Second, RWC_tree_ depends on the estimate of *W*, which is based on sapwood biomass in this study. If water exchange between sapwood and heartwood occurs (Treydte et al., 2021), the effective *W* may be larger than assumed here. Direct observations of tissue water content would therefore be particularly valuable for constraining physiologically realistic RWC_tree_ranges (e.g. Matheny et al., 2015; Martius et al., 2024).

Despite these uncertainties, the simulated outputs TWD_norm_ and RWC_tree_ provide physiologically interpretable diagnostic indicators of plant water status. According to Peters et al. (2025), TWD_norm_ = 1 marks the onset of drought stress and stomatal closure, while values around 1.6 indicate turgor loss in *Fagus sylvatica*. These thresholds were repeatedly exceeded at the Štítná site during dry years, indicating pronounced drought stress. Likewise, simulated RWC_tree_ declined to values around 0.86 in 2018, corresponding to a temporary depletion of roughly 14 % of the internal water pool. This degree of depletion is consistent with 2018 having been exceptionally warm and dry at the Štítná site (Nezval et al., 2025). In previous studies, comparable levels of depletion have been associated with hydraulic impairment in *Fagus sylvatica* seedlings (Rosner et al., 2019). Although mature trees are expected to tolerate somewhat greater depletion (Rosner et al., 2019), these values are consistent with pronounced physiological stress.

### 4.3 Boundary-case behavior under suppressed soil water recharge

The rainfall-suppression stress test represents a numerical boundary case in which the soil water pool was progressively depleted without replenishment (Fig. 7). In this setup, RWC_tree_ approached zero, corresponding to complete exhaustion of the modeled internal tree water storage. However, experimental studies have associated severe damage and mortality risk with RWC_tree_ values around 0.6 (Birami et al., 2018; Trueba et al., 2019; Sapes and Sala, 2021; Trifilò et al., 2023), and RWC_tree_ = 0 is unlikely to be reached under natural conditions. In LandscapeDNDC, no drought-induced tree mortality is currently implemented, and the depletion of RWC_tree_ towards zero is a direct consequence of residual *E* losses despite stomatal closure under sustained drought forcing. However, this progression likely overestimates water loss in natural systems, where severe tissue damage, mortality, and associated changes in residual water loss would alter the trajectory before complete depletion is reached. We therefore interpret the trajectory before RWC_tree_ ≈ 0.6 as the physiologically relevant part of the test, whereas the subsequent dynamics primarily reflect the numerical behavior of TWIST after conditions associated with severe damage or mortality have likely been exceeded. The stress test should therefore be interpreted as a numerical boundary case rather than a biologically realistic mortality trajectory.

These dynamics have direct implications for the interpretation of TWD_norm_. Because water depletion is limited to the available storage *W*, TWD plateaued once RWC_tree_ = 0 was reached. The maximum simulated TWD_norm_ values reached 26, but these primarily reflect continued water loss beyond physiologically meaningful conditions. A more physiologically interpretable reference point is therefore obtained at RWC_tree_ ≈ 0.6, where TWD_norm_ was approximately 10. This is higher than the reported value of ∼ 3 for drought-stressed mature *Fagus sylvatica* (Peters et al., 2025), but closer to the values reported for *Larix decidua* and *Pinus sylvestris* saplings at drought-induced tree death (8.4 and 12.0, respectively; TWD_norm_ time series shown in Fig. S5; Ziegler et al., 2024). However, such comparisons remain tentative, because published TWD_norm_ values near lethal dehydration are still scarce, the Peters et al. (2025) values refer to severe drought stress rather than mortality, and the values reported by Ziegler et al. (2024) were obtained for conifer saplings rather than mature beech trees.

### 4.4 Ways forward

A key way forward lies in applying TWIST to link expanding dendrometer observations with forest modeling frameworks. Because the module requires only *E*, *θ*_rel_, and an estimate of available internal water storage, it can be integrated into models that already simulate transpiration and soil moisture dynamics. The presented calibration workflow should further facilitate the derivation of site- and species-specific parameter sets where suitable evaluation data are available. With appropriate parameterization, the framework could in principle be applied across species, stand structures, and environmental conditions. This would allow TWIST to provide a diagnostic representation of internal tree water dynamics largely absent from current ecosystem models, enabling a direct link between such models and tree-level dendrometer drought-stress monitoring. A second priority for future work is to move beyond the diagnostic role of TWIST by implementing feedbacks on drought-sensitive processes in coupled models. For example, stomatal regulation (Peters et al., 2025) or minimum leaf conductance (Bartlett et al., 2012; Trueba et al., 2026) could be related to TWD_norm_ or RWC_tree_, allowing *E* to respond to progressive tree water depletion. Likewise, the close link of RWC_tree_ to cell turgor may provide a physiologically meaningful basis for representing drought-induced growth limitation and changes in carbon allocation (Potkay et al., 2022; Potkay and Feng, 2023). A further step could be to use RWC_tree_ thresholds or cumulative exposure to low-hydrated conditions to explore representations of progressive hydraulic impairment and mortality risk (Mantova et al., 2021; Mantova et al., 2022). However, such direct feedback is not yet implemented, and the corresponding threshold values require further evaluation before they can be applied more generally.

A further opportunity lies in testing whether RWC_tree_ can be related to remotely sensed indicators of canopy water status (Konings et al., 2019), thereby contributing to broader efforts to connect vegetation water content with ecosystem function (Binks et al., 2024). Because TWIST can be applied at stand scale while being constrained by tree-level dendrometer observations, it may provide a route for transferring physiological information from dendrometer measurements to broader ecosystem-scale assessments.

Combined datasets that link dendrometer records with coordinated ecosystem flux measurements, remote sensing products, soil moisture observations, sap flow, biomass estimates, or direct measurements of tissue water content would be particularly valuable for constraining TWIST parameters, testing transferability across sites and species, and developing feedback formulations that move TWIST beyond its current diagnostic role.

### 4.5 Conclusion

We presented TWIST, a diagnostic framework for translating imbalances between transpiration and water uptake into volume-based tree water deficit (TWD) dynamics. Despite its parsimonious structure, the module reproduced characteristic TWD patterns, including diurnal depletion–refilling cycles, incomplete nocturnal recovery, and progressive seasonal deficit accumulation. By generating a model state variable that can be compared with dendrometer-derived TWD, TWIST provides a practical interface between tree-level drought monitoring and process-based stand-level simulations. The derived relative tree water content (RWC_tree_) further expresses water deficits relative to an estimated internal water pool, which may also facilitate links to remotely sensed observations of canopy water status indicators. Together, TWD and RWC_tree_ provide complementary diagnostic indicators of internal tree hydration status, a component still often missing from current ecosystem models (Martinez-Vilalta et al., 2019). TWIST requires only a small number of inputs commonly available in process-based models, and the provided parameter calibration routine may support broader application. By making TWD available as a model-compatible diagnostic variable, TWIST creates a direct route for using expanding dendrometer datasets to evaluate and improve simulations of forest drought stress.

## 5 Acknowledgments

This study benefited from ongoing efforts to improve measurement standards and harmonize eddy covariance data processing within the FLUXNET and ICOS networks. In the preparation of this manuscript, the authors used ChatGPT (GPT-5.5, OpenAI) to support language refinement and code development for data analysis, plotting, and parameter calibration/evaluation. All AI-assisted output was reviewed, edited, and validated by the authors, who take full responsibility for the final content.

## 6 Authors’ Contributions

YZ developed and evaluated TWIST and wrote the manuscript draft. NKR and RG supported the conceptualization of the study and model development. PL and RG contributed to the LandscapeDNDC site setup and to the introduction and discussion. JK and LŠ provided observational data. MT contributed to the introduction. All authors discussed the results and revised the manuscript draft.

## 7 Supplementary Data

The following supplementary material is available for this article:

- **Supplemental Figure S1:** Evaluation of LandscapeDNDC-modeled Evapotranspiration
- **Supplemental Figure S2:** Evaluation of LandscapeDNDC-modeled Gross Primary Productivity
- **Supplemental Figure S3:** Evaluation of LandscapeDNDC-modeled Soil Water Content
- **Supplemental Figure S4:** Relationships between TWIST model parameters
- **Supplemental Figure S5:** Exemplary dendrometer-derived TWD data
- **Supplemental Table S1:** LandscapeDNDC parameters
- **Supplemental Table S2:** TWIST best-fit parameter values and distribution of acceptable solutions
- **Supplemental Table S3:** TWIST acceptable solutions parameter sets

## 10 Data and Code Availability

An R implementation of the TWIST module (version 2.0), including the core model functions, an example application, and workflows for parameter calibration and uncertainty analysis, together with all input data required to reproduce the TWIST simulations for the Štítná site (2018–2024), is openly available on GitHub (https://github.com/yanickziegler/TWIST) under the MIT License. An archived release with a DOI is additionally deposited at Zenodo (https://doi.org/10.5281/zenodo.19692465). The LandscapeDNDC ecosystem model used to generate the simulation drivers is available from the official project website (https://ldndc.imk-ifu.kit.edu/). The Štítná FLUXNET product was obtained from the ICOS Carbon Portal (https://hdl.handle.net/11676/sd5QIjbP_J9uxK7jRwbz5Bm1).

## References

Andriantelomanana, Tsiky, Thierry Améglio, Sylvain Delzon, Hervé Cochard, and Stephane Herbette (2024). “Unpacking the point of no return under drought in poplar: insight from stem diameter variation”. In: New Phytologist. doi: 10.1111/nph.19615.

Bartlett, Megan K., Christine Scoffoni, and Lawren Sack (2012). “The determinants of leaf turgor loss point and prediction of drought tolerance of species and biomes: a global meta-analysis”. en. In: Ecology Letters 15.5, pp. 393–405. doi: 10.1111/j.1461-0248.2012.01751.x.

Betsch, P., D. Bonal, N. Breda, P. Montpied, M. Peiffer, A. Tuzet, et al. (2011). “Drought effects on water relations in beech: The contribution of exchangeable water reservoirs”. en. In: Agricultural and Forest Meteorology 151.5, pp. 531–543. doi: 10.1016/j.agrformet.2010.12.008.

Binks, Oliver, Patrick Meir, Alexandra G. Konings, Lucas Cernusak, Bradley O. Christoffersen, William R. L. Anderegg, et al. (2024). “A Theoretical Framework to Quantify Ecosystem Pressure-Volume Relationships”. en. In: Global Change Biology 30.11, e17567. doi: 10.1111/gcb.17567.

Birami, Benjamin, Marielle Gattmann, Arnd G. Heyer, Rüdiger Grote, Almut Arneth, and Nadine K. Ruehr (2018). “Heat Waves Alter Carbon Allocation and Increase Mortality of Aleppo Pine Under Dry Conditions”. In: Frontiers in Forests and Global Change 1.November, pp. 1–17. doi: 10.3389/ffgc.2018.00008.

Blackman, Chris J., Danielle Creek, Chelsea Maier, Michael J. Aspinwall, John E. Drake, Sebastian Pfautsch, et al. (2019). “Drought response strategies and hydraulic traits contribute to mechanistic understanding of plant dry-down to hydraulic failure”. In: Tree Physiology 39.6, pp. 910–924. doi: 10.1093/treephys/tpz016.

Bose, Arun K., Sophia Etzold, Katrin Meusburger, Arthur Gessler, Andri Baltensweiler, Sabine Braun, et al. (2025). “Decreasing Stem Growth in Common European Tree Species Despite Earlier Growth Onset”. en. In: Global Change Biology 31.7, e70318. doi: 10.1111/gcb.70318.

Byrd, Richard H., Peihuang Lu, Jorge Nocedal, and Ciyou Zhu (1995). “A Limited Memory Algorithm for Bound Constrained Optimization”. en. In: SIAM Journal on Scientific Computing 16.5, pp. 1190–1208. doi: 10.1137/0916069.

Cade, Shirley M., Kevin C. Clemitshaw, Saúl Molina-Herrera, Rüdiger Grote, Edwin Haas, Matthew Wilkinson, et al. (2021). “Evaluation of LandscapeDNDC Model Predictions of CO2 and N2O Fluxes from an Oak Forest in SE England”. en. In: Forests 12.11, p. 1517. doi: 10.3390/f12111517.

Chen, Liangzhi, Philipp Brun, Pascal Buri, Simone Fatichi, Arthur Gessler, Michael James McCarthy, et al. (2025). “Global increase in the occurrence and impact of multiyear droughts”. en. In: Science 387.6731, pp. 278–284. doi: 10.1126/science.ado4245.

Cochard, Hervé, Eric Badel, Stéphane Herbette, Sylvain Delzon, Brendan Choat, and Steven Jansen (2013). “Methods for measuring plant vulnerability to cavitation: A critical review”. In: Journal of Experimental Botany 64.15, pp. 4779–4791. doi: 10.1093/jxb/ert193.

Cochard, Hervé, François Pimont, Julien Ruffault, and Nicolas Martin-StPaul (2021). “SurEau: a mechanistic model of plant water relations under extreme drought”. en. In: Annals of Forest Science 78.2, p. 55. doi: 10.1007/s13595-021-01067-y.

De Schepper, Veerle, Dagmar Van Dusschoten, Paul Copini, Siegfried Jahnke, and Kathy Steppe (2012). “MRI links stem water content to stem diameter variations in transpiring trees”. In: Journal of Experimental Botany 63.7, pp. 2645–2653. doi: 10.1093/jxb/err445.

De Swaef, Tom, Veerle De Schepper, Maurits W. Vandegehuchte, and Kathy Steppe (2015). “Stem diameter variations as a versatile research tool in ecophysiology”. In: Tree Physiology 35.10, pp. 1047–1061. doi: 10.1093/treephys/tpv080.

Dietrich, Lars, Roman Zweifel, and Ansgar Kahmen (2018). “Daily stem diameter variations can predict the canopy water status of mature temperate trees”. In: Tree Physiology 38.7, pp. 941–952. doi: 10.1093/treephys/tpy023.

Dietz, P. (1975). “Dichte und Rindengehalt von Industrieholz”. de. In: Holz als Roh- und Werkstoff 33.4, pp. 135–141. doi: 10.1007/BF02611237.

Dirnböck, T., D. Kraus, R. Grote, S. Klatt, J. Kobler, A. Schindlbacher, et al. (2020). “Substantial understory contribution to the C sink of a European temperate mountain forest landscape”. en. In: Landscape Ecology 35.2, pp. 483–499. doi: 10.1007/s10980-019-00960-2.

Duursma, Remko A., Christopher J. Blackman, Rosana Lopéz, Nicolas K. Martin-StPaul, Hervé Cochard, and Belinda E. Medlyn (2019). “On the minimum leaf conductance: its role in models of plant water use, and ecological and environmental controls”. en. In: New Phytologist 221.2, pp. 693–705. doi: 10.1111/nph.15395.

Eller, Cleiton B., Lucy Rowland, Maurizio Mencuccini, Teresa Rosas, Karina Williams, Anna Harper, et al. (2020). “Stomatal optimization based on xylem hydraulics (SOX) improves land surface model simulation of vegetation responses to climate”. In: New Phytologist 226.6, pp. 1622–1637. doi: 10.1111/nph.16419.

Feng, Feng, Yael Wagner, Tamir Klein, and Uri Hochberg (2023). “Xylem resistance to cavitation increases during summer in Pinus halepensis”. en. In: Plant, Cell & Environment 46.6, pp. 1849–1859. doi: 10.1111/pce.14573.

Gazol, Antonio, Manuel Pizarro, William M. Hammond, Craig D. Allen, and J. Julio Camarero (2025). “Droughts preceding tree mortality events have increased in duration and intensity, especially in dry biomes”. en. In: Nature Communications 16.1, p. 5779. doi: 10.1038/s41467-025-60856-5.

Gleason, Sean M., Chris J. Blackman, Alicia M. Cook, Claire A. Laws, and Mark Westoby (2014). “Whole-plant capacitance, embolism resistance and slow transpiration rates all contribute to longer desiccation times in woody angiosperms from arid and wet habitats”. In: Tree Physiology 34.3, pp. 275–284. doi: 10.1093/treephys/tpu001.

Grote, R., J. Korhonen, and I. Mammarella (2011a). “Challenges for evaluating process-based models of gas exchange at forest sites with fetches of various species”. In: Investigacion Agraria Sistemas y Recursos Forestales 20.3, pp. 389–406. doi: 10.5424/fs/20112003-11084.

Grote, Rüdiger, Ralf Kiese, Thomas Grünwald, Jean Marc Ourcival, and André Granier (2011b). “Modelling forest carbon balances considering tree mortality and removal”. In: Agricultural and Forest Meteorology 151.2, pp. 179–190. doi: 10.1016/j.agrformet.2010.10.002.

Haas, Edwin, Steffen Klatt, Alexander Fröhlich, Philipp Kraft, Christian Werner, Ralf Kiese, et al. (2013). “LandscapeDNDC: A process model for simulation of biosphere-atmosphere-hydrosphere exchange processes at site and regional scale”. In: Landscape Ecology 28.4, pp. 615–636. doi: 10.1007/s10980-012-9772-x.

Hartmann, Henrik, Catarina F. Moura, William R. L. Anderegg, Nadine K. Ruehr, Yann Salmon, Craig D. Allen, et al. (2018). “Research frontiers for improving our understanding of drought-induced tree and forest mortality”. en. In: New Phytologist 218.1, pp. 15–28. doi: 10.1111/nph.15048.

International Tree Mortality Network (2025). “Towards a global understanding of tree mortality”. en. In: New Phytologist 245.6, pp. 2377–2392. doi: 10.1111/nph.20407.

Klein, T., S. Cohen, I. Paudel, Y. Preisler, E. Rotenberg, and D. Yakir (2016). “Diurnal dynamics of water transport, storage and hydraulic conductivity in pine trees under seasonal drought”. en. In: iForest - Biogeosciences and Forestry 9.5, p. 710. doi: 10.3832/ifor2046-009.

Klein, Tamir, Eyal Rotenberg, Ella Cohen-Hilaleh, Naama Raz-Yaseef, Fyodor Tatarinov, Yakir Preisler, et al. (2014). “Quantifying transpirable soil water and its relations to tree water use dynamics in a water-limited pine forest”. en. In: Ecohydrology 7.2, pp. 409–419. doi: 10.1002/eco.1360.

Konings, Alexandra G., Krishna Rao, and Susan C. Steele-Dunne (2019). “Macro to micro: microwave remote sensing of plant water content for physiology and ecology”. en. In: New Phytologist 223.3, pp. 1166–1172. doi: 10.1111/nph.15808.

Kraus, David, Christian Werner, Baldur Janz, Steffen Klatt, Björn Ole Sander, Reiner Wassmann, et al. (2022). “Greenhouse Gas Mitigation Potential of Alternate Wetting and Drying for Rice Production at National Scale—A Modeling Case Study for the Philippines”. en. In: Journal of Geophysical Research: Biogeosciences 127.5, e2022JG006848. doi: 10.1029/2022JG006848.

Köcher, Paul, Viviana Horna, and Christoph Leuschner (2013). “Stem water storage in five coexisting temperate broad-leaved tree species: significance, temporal dynamics and dependence on tree functional traits”. In: Tree Physiology 33.8, pp. 817–832. doi: 10.1093/treephys/tpt055.

Lawlor, D. W. and G. Cornic (2002). “Photosynthetic carbon assimilation and associated metabolism in relation to water deficits in higher plants”. en. In: Plant, Cell & Environment 25.2, pp. 275–294. doi: 10.1046/j.0016-8025.2001.00814.x.

Li, Shan, Marion Feifel, Zohreh Karimi, Bernhard Schuldt, Brendan Choat, and Steven Jansen (2016). “Leaf gas exchange performance and the lethal water potential of five European species during drought”. In: Tree Physiology 36.2, pp. 179–192. doi: 10.1093/treephys/tpv117.

Mantova, Marylou, Stéphane Herbette, Hervé Cochard, and José M. Torres-Ruiz (2022). “Hydraulic failure and tree mortality: from correlation to causation”. In: Trends in Plant Science 27.4, pp. 335–345. doi: 10.1016/j.tplants.2021.10.003.

Mantova, Marylou, Paulo E. Menezes-Silva, Eric Badel, Hervé Cochard, and José M. Torres-Ruiz (2021). “The interplay of hydraulic failure and cell vitality explains tree capacity to recover from drought”. In: Physiologia Plantarum 172.1, pp. 247–257. doi: 10.1111/ppl.13331.

Martinez-Vilalta, Jordi, William R. L. Anderegg, Gerard Sapes, and Anna Sala (2019). “Greater focus on water pools may improve our ability to understand and anticipate drought-induced mortality in plants”. en. In: New Phytologist 223.1, pp. 22–32. doi: 10.1111/nph.15644.

Martius, Lion R, Maurizio Mencuccini, Paulo R L Bittencourt, Moisés Moraes Alves, Oliver Binks, Pablo Sanchez-Martinez, et al. (2024). “Towards accurate monitoring of water content in woody tissue across tropical forests and other biomes”. In: Tree Physiology 44.8, tpae076. doi: 10.1093/treephys/tpae076.

Matheny, Ashley M., Gil Bohrer, Steven R. Garrity, Timothy H. Morin, Cecil J. Howard, and Christoph S. Vogel (2015). “Observations of stem water storage in trees of opposing hydraulic strategies”. en. In: Ecosphere 6.9, art165. doi: 10.1890/ES15-00170.1.

McDowell, Nate G., Gerard Sapes, Alexandria Pivovaroff, Henry D. Adams, Craig D. Allen, William R.L. Anderegg, et al. (2022). “Mechanisms of woody-plant mortality under rising drought, CO2 and vapour pressure deficit”. In: Nature Reviews Earth and Environment 3.5, pp. 294–308. doi: 10.1038/s43017-022-00272-1.

McGloin, Ryan, Ladislav Šigut, Milan Fischer, Lenka Foltýnová, Shilpi Chawla, Miroslav Trnka, et al. (2019). “Available Energy Partitioning During Drought at Two Norway Spruce Forests and a European Beech Forest in Central Europe”. en. In: Journal of Geophysical Research: Atmospheres 124.7, pp. 3726–3742. doi: 10.1029/2018JD029490.

Meyer, Benjamin F., João P. Darela-Filho, Konstantin Gregor, Allan Buras, Qiao-Lin Gu, Andreas Krause, et al. (2025). “Simulating the drought response of European tree species with the dynamic vegetation model LPJ-GUESS (v4.1, 97c552c5)”. English. In: Geoscientific Model Development 18.14, pp. 4643–4666. doi: 10.5194/gmd-18-4643-2025.

Missik, Justine E. C., Gil Bohrer, Madeline E. Scyphers, Ashley M. Matheny, Ana Maria Restrepo Acevedo, Marcela Silva, et al. (2025). “Using a Plant Hydrodynamic Model, FETCH4, to Supplement Measurements and Characterize Hydraulic Traits in a Mixed Temperate Forest”. en. In: Journal of Geophysical Research: Biogeosciences 130.4, e2024JG008198. doi: 10.1029/2024JG008198.

Molina-Herrera, Saúl, Edwin Haas, Steffen Klatt, David Kraus, Jürgen Augustin, Vincenzo Magliulo, et al. (2016). “A modeling study on mitigation of N2O emissions and NO3 leaching at different agricultural sites across Europe using LandscapeDNDC”. In: Science of The Total Environment 553, pp. 128–140. doi: 10.1016/j.scitotenv.2015.12.099.

Müller, Lena M. and Michael Bahn (2022). “Drought legacies and ecosystem responses to subsequent drought”. In: Global Change Biology 28.17, pp. 5086–5103. doi: 10.1111/gcb.16270.

Nadal-Sala, Daniel, Rüdiger Grote, David Kraus, Uri Hochberg, Tamir Klein, Yael Wagner, et al. (2024). “Integration of tree hydraulic processes and functional impairment to capture the drought resilience of a semiarid pine forest”. en. In: Biogeosciences 21.12, pp. 2973–2994. doi: 10.5194/bg-21-2973-2024.

Nezval, Ondřej, Lenka Foltýnová, Marek Fajstavr, Jan Krejza, Ladislav Šigut, Jan Světlík, et al. (2025). “Temperature-Driven onset and light quality-linked senescence in Fagus sylvatica phenology”. en. In: Agricultural and Forest Meteorology 375, p. 110834. doi: 10.1016/j.agrformet.2025.110834.

Novick, Kimberly A., Darren L. Ficklin, Dennis Baldocchi, Kenneth J. Davis, Teamrat A. Ghezzehei, Alexandra G. Konings, et al. (2022). “Confronting the water potential information gap”. In: Nature Geoscience 15.3, pp. 158–164. doi: 10.1038/s41561-022-00909-2.

Oberhuber, Walter, Andreas Gruber, and Gerhard Wieser (2023). “Seasonal and daily xylem radius variations in scots pine are closely linked to environmental factors affecting transpiration”. In: Biology 12.9. doi: 10.3390/biology12091251.

Pastorello, Gilberto, Carlo Trotta, Eleonora Canfora, Housen Chu, Danielle Christianson, You-Wei Cheah, et al. (2020). “The FLUXNET2015 dataset and the ONEFlux processing pipeline for eddy covariance data”. en. In: Scientific Data 7.1, p. 225. doi: 10.1038/s41597-020-0534-3.

Peramaki, M., E. Nikinmaa, S. Sevanto, H. Ilvesniemi, E. Siivola, P. Hari, et al. (2001). “Tree stem diameter variations and transpiration in Scots pine: an analysis using a dynamic sap flow model”. en. In: Tree Physiology 21.12-13, pp. 889–897. doi: 10.1093/treephys/21.12-13.889.

Peramaki, M., T. Vesala, and E. Nikinmaa (2005). “Modeling the dynamics of pressure propagation and diameter variation in tree sapwood”. en. In: Tree Physiology 25.9, pp. 1091–1099. doi: 10.1093/treephys/25.9.1091.

Peters, Richard L., David Basler, Roman Zweifel, David N. Steger, Tobias Zhorzel, Cedric Zahnd, et al. (2025). “Normalized tree water deficit: an automated dendrometer signal to quantify drought stress in trees”. en. In: New Phytologist, nph.70266. doi: 10.1111/nph.70266.

Peters, Richard L., Kathy Steppe, Christoforos Pappas, Roman Zweifel, Flurin Babst, Lars Dietrich, et al. (2023). “Daytime stomatal regulation in mature temperate trees prioritizes stem rehydration at night”. In: New Phytologist. doi: 10.1111/nph.18964.

Potkay, Aaron and Xue Feng (2023). “Do stomata optimize turgor-driven growth? A new framework for integrating stomata response with whole-plant hydraulics and carbon balance”. en. In: New Phytologist 238.2, pp. 506–528. doi: 10.1111/nph.18620.

Potkay, Aaron, Teemu Hölttä, Anna T Trugman, and Ying Fan (2022). “Turgor-limited predictions of tree growth, height and metabolic scaling over tree lifespans”. en. In: Tree Physiology 42.2. Ed. by Michael Ryan, pp. 229–252. doi: 10.1093/treephys/tpab094.

Preisler, Yakir, Teemu Hölttä, José M Grünzweig, Itay Oz, Fedor Tatarinov, Nadine K Ruehr, et al. (2022). “The importance of tree internal water storage under drought conditions”. en. In: Tree Physiology 42.4. Ed. by Ram Oren, pp. 771–783. doi: 10.1093/treephys/tpab144.

Preisler, Yakir, Fyodor Tatarinov, José M. Grünzweig, Didier Bert, Jérôme Ogée, Lisa Wingate, et al. (2019). “Mortality versus survival in drought-affected Aleppo pine forest depends on the extent of rock cover and soil stoniness”. en. In: Functional Ecology 33.5, pp. 901–912. doi: 10.1111/1365-2435.13302.

R Core Team (2022). R: A Language and Environment for Statistical Computing.

Restrepo-Acevedo, Ana Maria, Jessica Guo, Kimberly Novick, Vincent Humphrey, Roman Zweifel, Alexandra G. Konings, et al. (2026). “Continuous monitoring of plant water potential: sensor-based approaches and best practices”. en. In: New Phytologist, nph.71194. doi: 10.1111/nph.71194.

Rosner, Sabine, Berthold Heinze, Tadeja Savi, and Guillermina Dalla-Salda (2019). “Prediction of hydraulic conductivity loss from relative water loss: new insights into water storage of tree stems and branches”. In: Physiologia Plantarum 165.4, pp. 843–854. doi: 10.1111/ppl.12790.

Ruffault, Julien, François Pimont, Hervé Cochard, Jean-Luc Dupuy, and Nicolas Martin-StPaul (2022). “SurEau-Ecos v2.0: a trait-based plant hydraulics model for simulations of plant water status and drought-induced mortality at the ecosystem level”. en. In: Geoscientific Model Development 15.14, pp. 5593–5626. doi: 10.5194/gmd-15-5593-2022.

Salomón, Roberto L., Jean Marc Limousin, Jean Marc Ourcival, Jesús Rodríguez-Calcerrada, and Kathy Steppe (2017). “Stem hydraulic capacitance decreases with drought stress: implications for modelling tree hydraulics in the Mediterranean oak Quercus ilex”. In: Plant Cell and Environment 40.8, pp. 1379–1391. doi: 10.1111/pce.12928.

Sapes, Gerard and Anna Sala (2021). “Relative water content consistently predicts drought mortality risk in seedling populations with different morphology, physiology and times to death”. en. In: Plant, Cell & Environment 44.10, pp. 3322–3335. doi: 10.1111/pce.14149.

Senf, Cornelius, Allan Buras, Christian S. Zang, Anja Rammig, and Rupert Seidl (2020). “Excess forest mortality is consistently linked to drought across Europe”. In: Nature Communications 11.1, pp. 1–8. doi: 10.1038/s41467-020-19924-1.

Sevanto, Sanna (2001). “Xylem diameter changes as an indicator of stand-level evapo-transpiration”. en. In: BOREAL ENVIRONMENT RESEARCH 6.

Sevanto, Sanna (2025). “Dendrometers—what are they good for?” In: Tree Physiology 45.4, tpaf035. doi: 10.1093/treephys/tpaf035.

Sifounakis, Odysseas, Edwin Haas, Klaus Butterbach-Bahl, and Maria P. Papadopoulou (2024). “Regional assessment and uncertainty analysis of carbon and nitrogen balances at cropland scale using the ecosystem model LandscapeDNDC”. English. In: Biogeosciences 21.6, pp. 1563–1581. doi: 10.5194/bg-21-1563-2024.

Steppe, Kathy, Dirk J.W. De Pauw, Raoul Lemeur, and Peter A. Vanrolleghem (2006). “A mathematical model linking tree sap flow dynamics to daily stem diameter fluctuations and radial stem growth”. In: Tree Physiology 26.3, pp. 257–273. doi: 10.1093/treephys/26.3.257.

Sánchez-Costa, Elisenda, Rafael Poyatos, and Santiago Sabaté (2015). “Contrasting growth and water use strategies in four co-occurring Mediterranean tree species revealed by concurrent measurements of sap flow and stem diameter variations”. In: Agricultural and Forest Meteorology 207, pp. 24–37. doi: 10.1016/j.agrformet.2015.03.012.

Tai, Xiaonan, D. Scott Mackay, John S. Sperry, Paul Brooks, William R. L. Anderegg, Lawrence B. Flanagan, et al. (2018). “Distributed Plant Hydraulic and Hydrological Modeling to Understand the Susceptibility of Riparian Woodland Trees to Drought-Induced Mortality”. en. In: Water Resources Research 54.7, pp. 4901–4915. doi: 10.1029/2018WR022801.

Treydte, Kerstin, Marco M Lehmann, Tomasz Wyczesany, and Sebastian Pfautsch (2021). “Radial and axial water movement in adult trees recorded by stable isotope tracing”. en. In: Tree Physiology 41.12. Ed. by Lucas Cernusak, pp. 2248–2261. doi: 10.1093/treephys/tpab080.

Trifilò, Patrizia, Elisa Abate, Francesco Petruzzellis, Maria Azzarà, and Andrea Nardini (2023). “Critical water contents at leaf, stem and root level leading to irreversible drought-induced damage in two woody and one herbaceous species”. en. In: Plant, Cell & Environment 46.1, pp. 119–132. doi: 10.1111/pce.14469.

Trueba, Santiago, Régis Burlett, Xavier Bouteiller, Thibaut Burneau, Guillaume Forget, Maximilien Larter, et al. (2026). “Ecological drivers and phylogenetic patterns of leaf minimum conductance variability in vascular plants”. en. In: New Phytologist n/a.n/a. doi: 10.1111/nph.71166.

Trueba, Santiago, Ruihua Pan, Christine Scoffoni, Grace P. John, Stephen D. Davis, and Lawren Sack (2019). “Thresholds for leaf damage due to dehydration: declines of hydraulic function, stomatal conductance and cellular integrity precede those for photochemistry”. en. In: New Phytologist 223.1, pp. 134–149. doi: 10.1111/nph.15779.

Verbeeck, Hans, Kathy Steppe, Nadja Nadezhdina, Maarten Op de Beeck, Gaby Deckmyn, Linda Meiresonne, et al. (2007). “Stored water use and transpiration in Scots pine: a modeling analysis with ANAFORE”. In: Tree Physiology 27.12, pp. 1671–1685.

Wagenführ, R (2007). Holzatlas. Vol. 6, neu bearbeitete und erweiterte Auflage. Leipzig, Germany: Fachbuchverlag Leipzig.

Xu, Chonggang, Bradley Christoffersen, Zachary Robbins, Ryan Knox, Rosie A. Fisher, Rutuja Chitra-Tarak, et al. (2023). “Quantification of hydraulic trait control on plant hydrodynamics and risk of hydraulic failure within a demographic structured vegetation model in a tropical forest (FATES–HYDRO V1.0)”. English. In: Geoscientific Model Development 16.21, pp. 6267–6283. doi: 10.5194/gmd-16-6267-2023.

Xu, Xiangtao, David Medvigy, Jennifer S. Powers, Justin M. Becknell, and Kaiyu Guan (2016). “Diversity in plant hydraulic traits explains seasonal and inter-annual variations of vegetation dynamics in seasonally dry tropical forests”. en. In: New Phytologist 212.1, pp. 80–95. doi: 10.1111/nph.14009.

Yao, Yitong, Emilie Joetzjer, Philippe Ciais, Nicolas Viovy, Fabio Cresto Aleina, Jerome Chave, et al. (2022). “Forest fluxes and mortality response to drought: model description (ORCHIDEE-CAN-NHA r7236) and evaluation at the Caxiuanã drought experiment”. English. In: Geoscientific Model Development 15.20, pp. 7809–7833. doi: 10.5194/gmd-15-7809-2022.

Ziegler, Yanick, Rüdiger Grote, Franklin Alongi, Timo Knüver, and Nadine K. Ruehr (2024). “Capturing drought stress signals: The potential of dendrometers for monitoring tree water status.” In: Tree Physiology. doi: 10.1093/treephys/tpae140.

Zweifel, R. and R. Häsler (2000). “Frost-induced reversible shrinkage of bark of mature subalpine conifers”. en. In: Agricultural and Forest Meteorology 102.4, pp. 213–222. doi: 10.1016/S0168-1923(00)00135-0.

Zweifel, R. and R. Häsler (2001). “Dynamics of water storage in mature subalpine Picea abies: temporal and spatial patterns of change in stem radius”. In: Tree Physiology.

Zweifel, R., H. Item, and R. Häsler (2001). “Link between diurnal stem radius changes and tree water relations”. In: Tree Physiology 21.12–13, pp. 869–877. doi: 10.1093/treephys/21.12-13.869.

Zweifel, R., L. Zimmermann, and D. M. Newbery (2005). “Modeling tree water deficit from microclimate: An approach to quantifying drought stress”. In: Tree Physiology 25.2, pp. 147–156. doi: 10.1093/treephys/25.2.147.

Zweifel, Roman, Sophia Etzold, David Basler, Reinhard Bischoff, Sabine Braun, Nina Buchmann, et al. (2021a). “TreeNet–The Biological Drought and Growth Indicator Network”. In: Frontiers in Forests and Global Change 4. doi: 10.3389/ffgc.2021.776905.

Zweifel, Roman, Matthias Haeni, Nina Buchmann, and Werner Eugster (2016). “Are trees able to grow in periods of stem shrinkage?” In: New Phytologist 211.3, pp. 839–849. doi: 10.1111/nph.13995.

Zweifel, Roman, Frank Sterck, Sabine Braun, Nina Buchmann, Werner Eugster, Arthur Gessler, et al. (2021b). “Why trees grow at night”. In: New Phytologist 231.6, pp. 2174–2185. doi: 10.1111/nph.17552.

Čermák, Jan, Jiří Kučera, William L. Bauerle, Nathan Phillips, and Thomas M. Hinckley (2007). “Tree water storage and its diurnal dynamics related to sap flow and changes in stem volume in old-growth Douglas-fir trees”. In: Tree Physiology 27.2, pp. 181–198. doi: 10.1093/treephys/27.2.181.

