## Supplementary material for "TWIST: A diagnostic framework for representing tree water deficit dynamics in process-based forest models"

The following supplementary material is available for this article:

- **Supplemental Figure S1:** Evaluation of LandscapeDNDC-modeled Evapotranspiration
- **Supplemental Figure S2:** Evaluation of LandscapeDNDC-modeled Gross Primary Productivity
- **Supplemental Figure S3:** Evaluation of LandscapeDNDC-modeled Soil Water Content
- **Supplemental Figure S4:** Relationships between TWIST model parameters
- **Supplemental Figure S5:** Exemplary dendrometer-derived TWD data
- **Supplemental Table S1:** LandscapeDNDC parameters
- **Supplemental Table S2:** TWIST best-fit parameter values and distribution of acceptable solutions
- **Supplemental Table S3:** TWIST acceptable solutions parameter sets

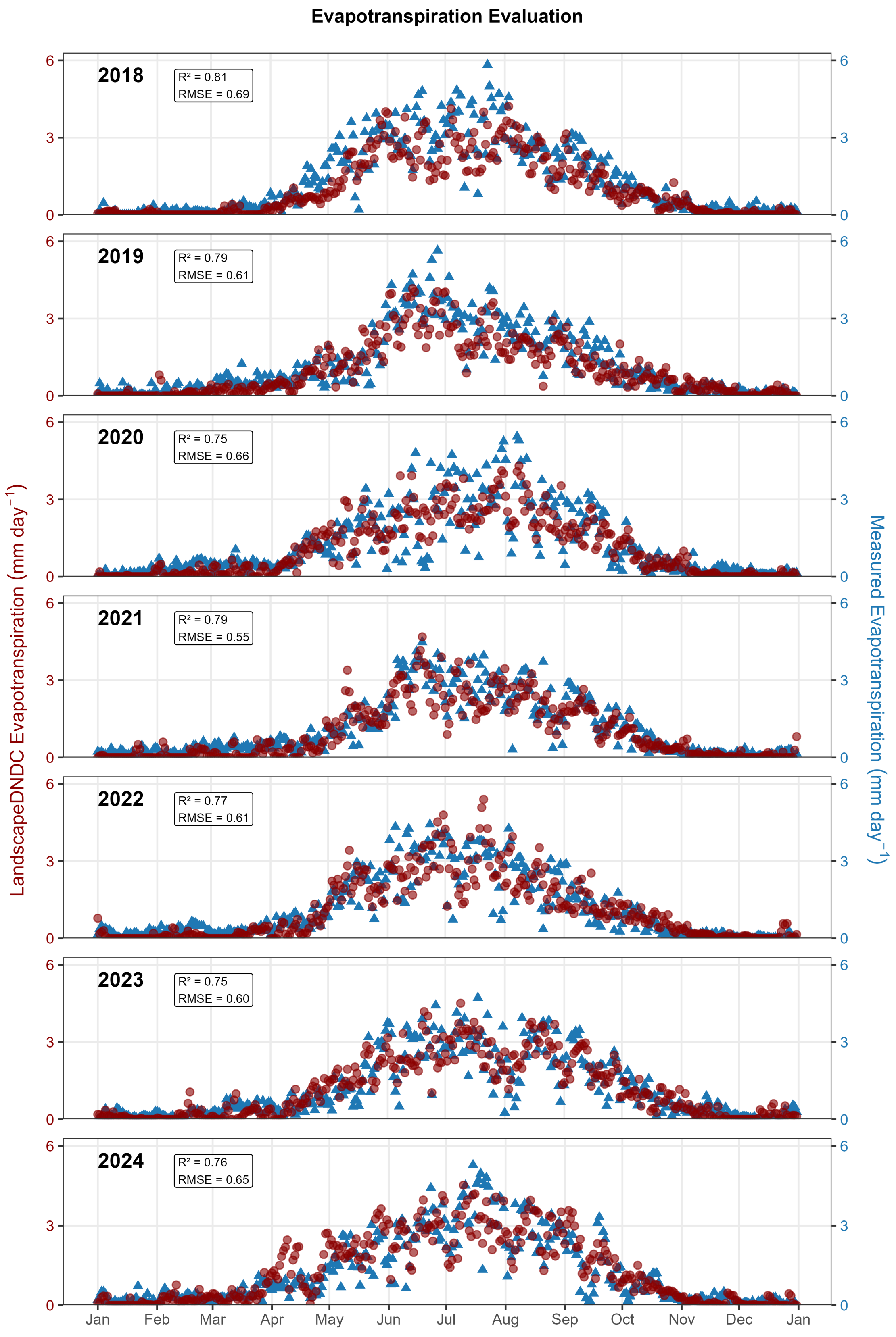


Figure S1: Evaluation of LandscapeDNDC-modeled Evapotranspiration (red) against observations (blue) at the Štítná forest site for the years 2018–2024, with statistical metrics ($R^2$, RMSE) quantifying model–data agreement.

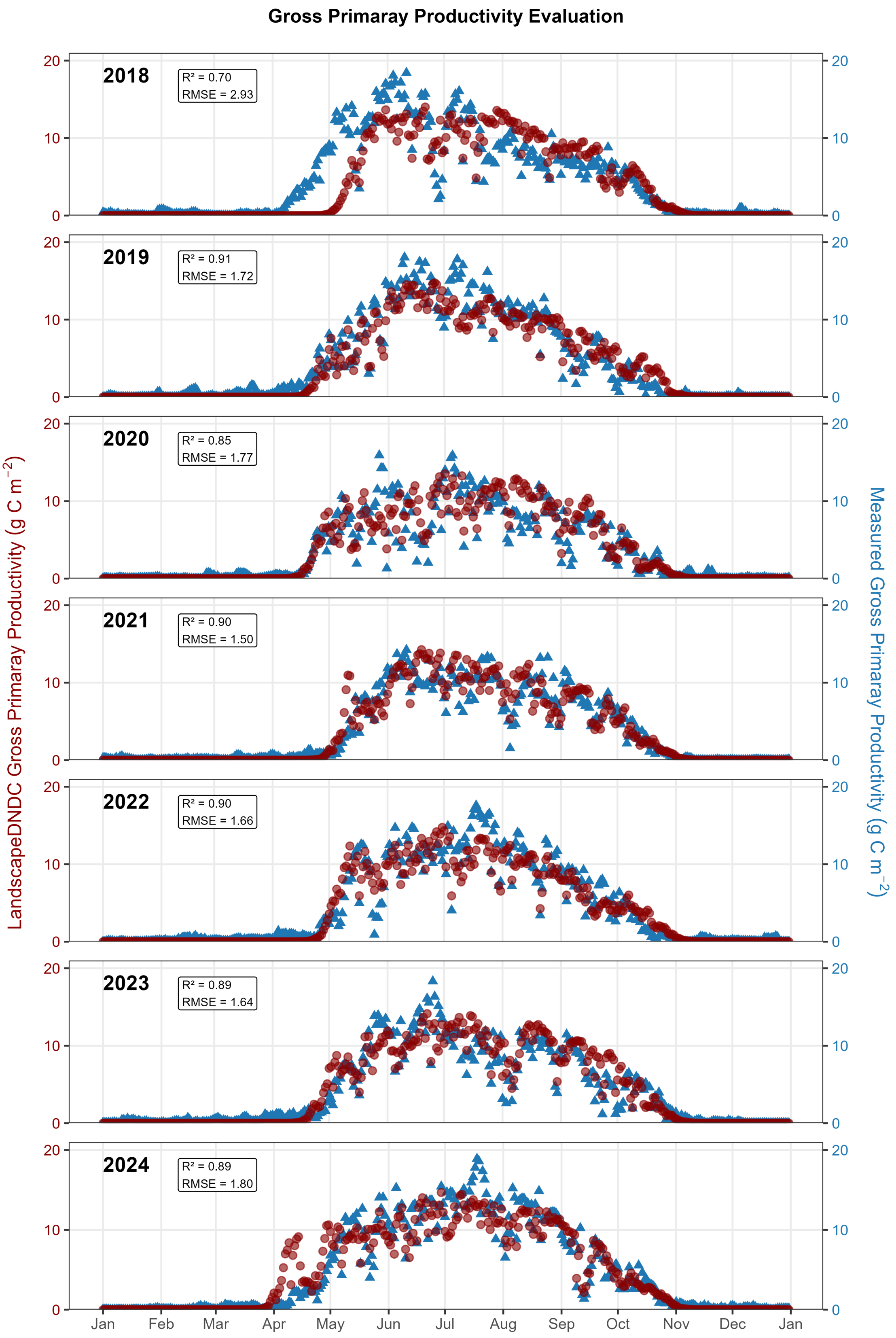


Figure S2: Evaluation of LandscapeDNDC-modeled Gross Primary Productivity (red) against observations (blue) at the Štítná forest site for the years 2018–2024, with statistical metrics ($R^2$, RMSE) quantifying model–data agreement.

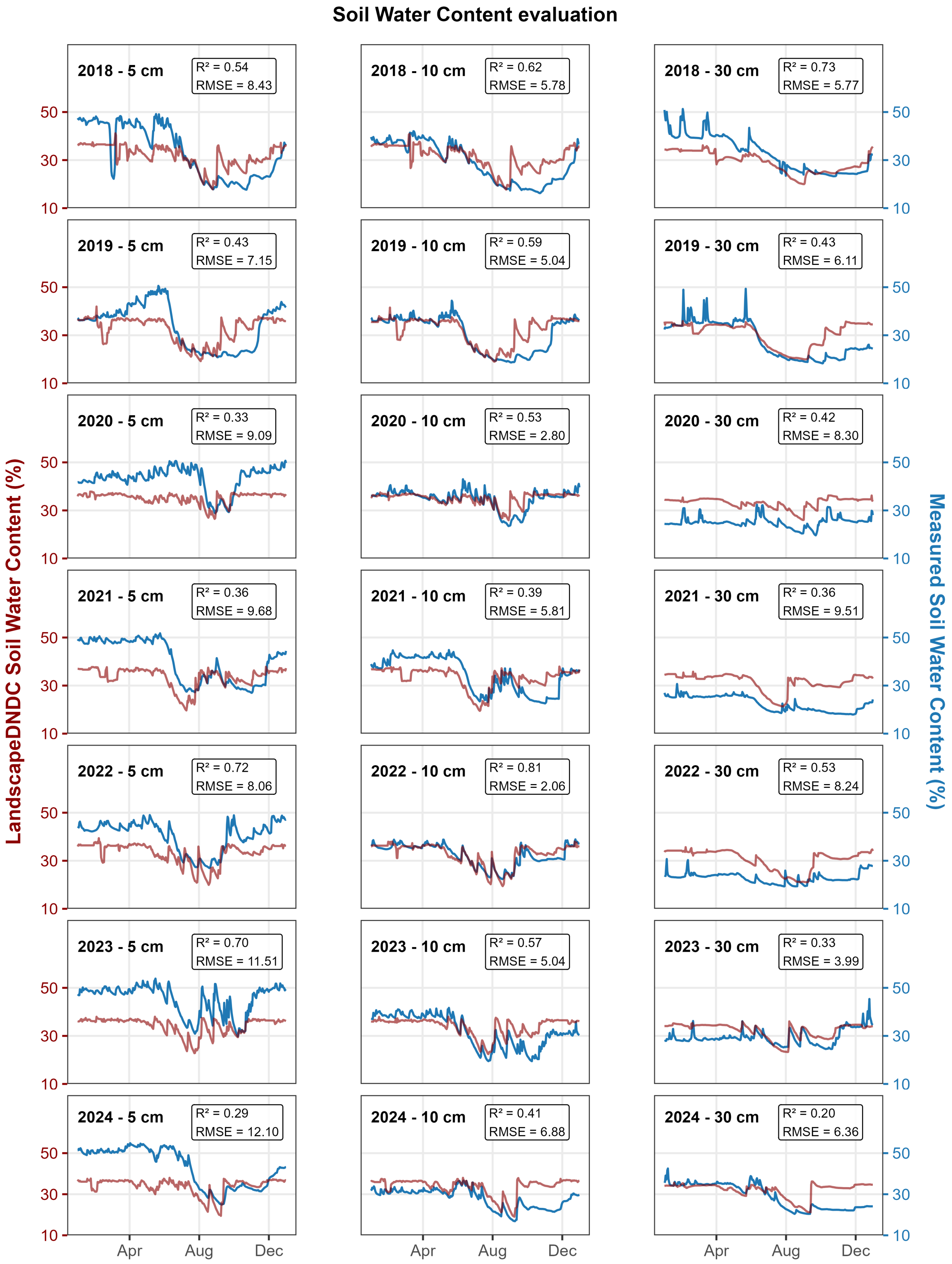


Figure S3: Evaluation of LandscapeDNDC-modeled soil water content (red) against measured (blue) at the Štítná forest site for the years 2018–2024 in three different depths, with statistical metrics ($R^2$, RMSE) quantifying model–data agreement.

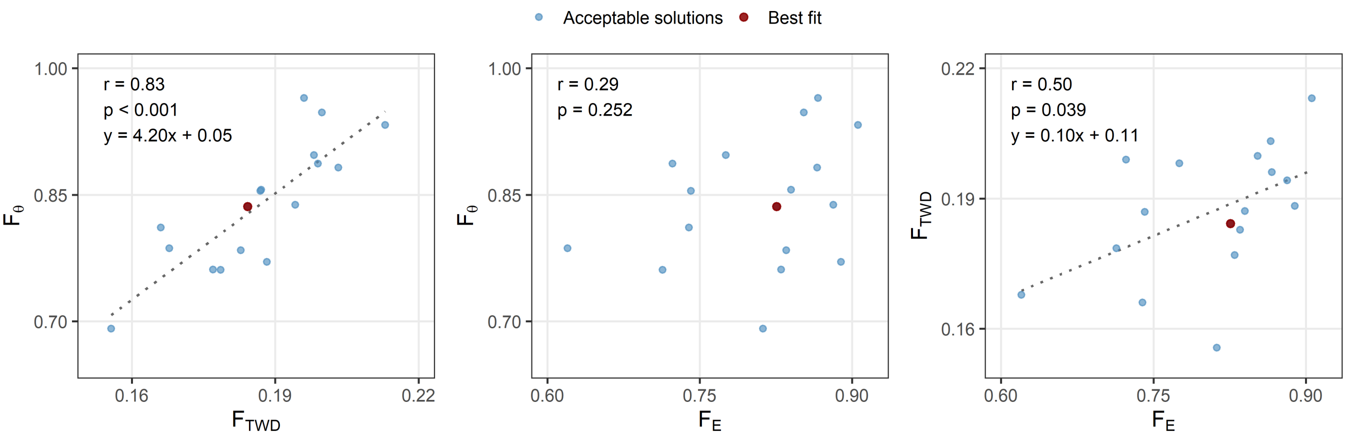


Figure S4: Relationships between TWIST model parameters based on the subset of acceptable solutions (defined as parameter sets with an objective value within 5% of the best-fit objective value; n=17). Scatterplots show pairwise relationships between the parameters $F_E$, $F_{\mathrm{TWD}}$ and $F_\theta$. Blue points represent acceptable parameter combinations, and the red point indicates the best-fit solution. The annotated values denote the Pearson correlation coefficient (r) between parameter pairs across all acceptable solutions, indicating the strength and direction of their linear dependency. A linear regression fit (dashed line) is only shown if a significant relationship was evident (p<0.05).

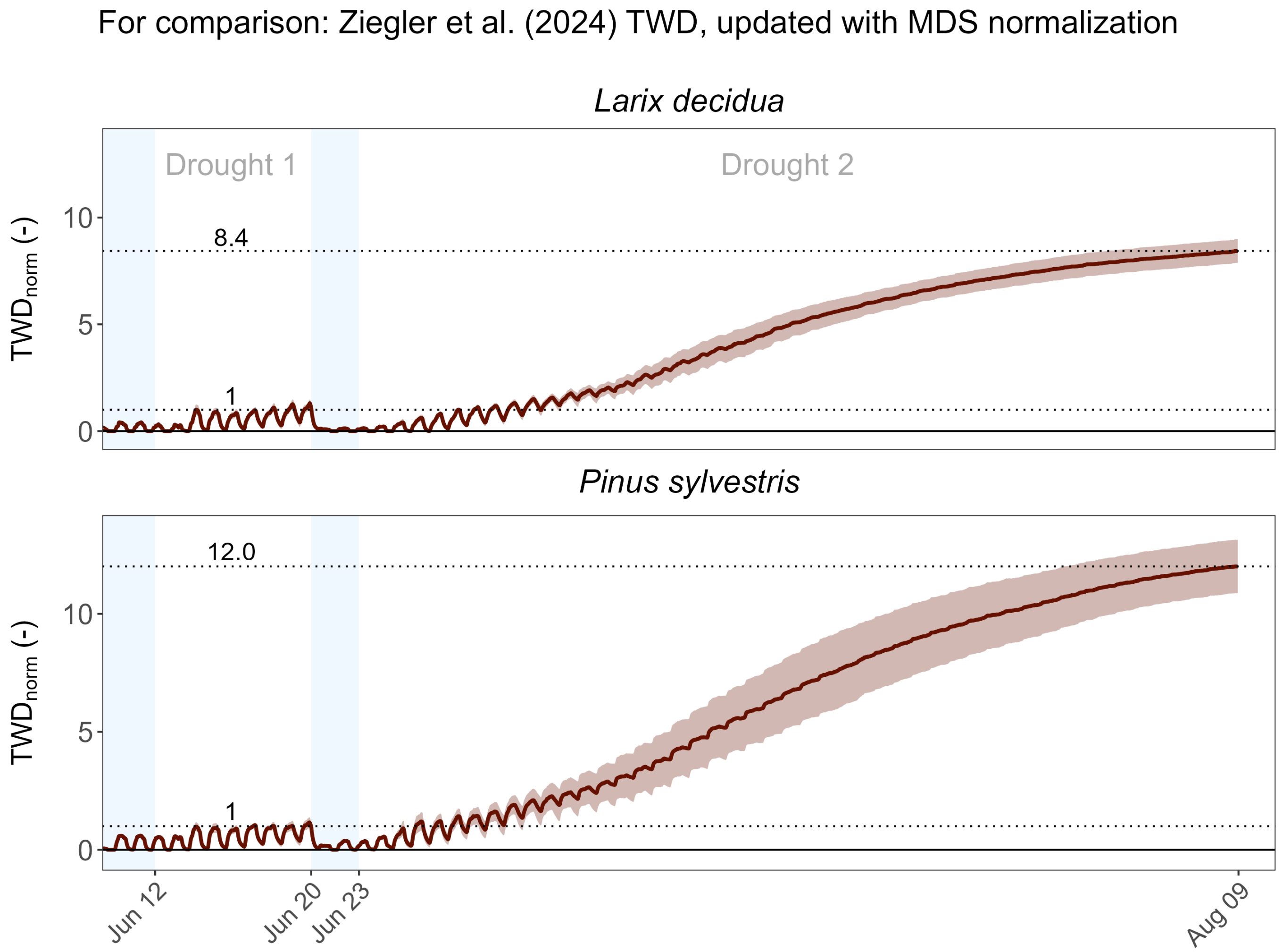


Figure S5: Dendrometer-derived TWD development under lethal dehydration in a greenhouse experiment covering two species, adapted from Ziegler et al. (2024). In contrast to the original paper, the normalization procedure was changed to MDS normalization (Peters et al., 2025) applied to the treatment mean timeseries to allow comparison with the current study.

Table S1: Selected LandscapeDNDC parameters required for transpiration demand and stomatal conductance calculations at the Štítná *Fagus sylvatica* site.

| **Parameter** | **Unit** | **Value** | **Description** |
|---|---|---|---|
| RPMIN | mmol $m^{-1}$ $m^2$LA s MPa | 1.4 | Minimum whole-plant resistance |
| GSMIN | mmol $H_2O$ $m^{-2}$ $s^{-1}$ | 4.3 | Minimum leaf conductance |
| GSMAX | mmol $H_2O$ $m^{-2}$ $s^{-1}$ | 81.5 | Maximum leaf conductance |
| A_EXP | – | 6.2 | Curve parameter for non-stomatal-limitation effect on photosynthesis |
| A_REF | MPa | -4.8 | Reference $\psi$ for non-stomatal-limitation effect on photosynthesis |
| CWP_REF | mmol $MPa^{-1}$ $m^{-2}$ $s^{-1}$ | 10 | Specific xylem conductance |
| PSI_EXP | – | 6.5 | Curve parameter for $\psi$ impact on conductance |
| PSI_REF | MPa | -3.6 | Reference $\psi$ for conductance vulnerability curve |
| MFOLOPT | kgDW $m^{-2}$ground | 0.25 | Foliage biomass for mature stands |

Table S2: TWIST best-fit parameter values and distribution of acceptable solutions. Acceptable solutions ($n = 17$) are defined based on the 2018 calibration period as parameter sets yielding calibration performance within 5% of the best-fit solution (see Methods). Reported are the best-fit value, median (q50), interquartile range (q25–q75), and full acceptable range (min–max) for each parameter. Spearman rank correlation coefficients ($\rho$), calculated across the full 2018–2024 time series, indicate the sensitivity of model performance to parameter variation.

| **Parameter** | **Best fit** | **min** | **q25** | **median** | **q75** | **max** | **Spearman $\rho$** |
|---|---|---|---|---|---|---|---|
| $F_E$ | 0.83 | 0.62 | 0.74 | 0.83 | 0.87 | 0.91 | -0.05 |
| $F_{\mathrm{TWD}}$ | 0.19 | 0.16 | 0.18 | 0.19 | 0.20 | 0.21 | 0.29 |
| $F_\theta$ | 0.84 | 0.69 | 0.78 | 0.84 | 0.89 | 0.96 | -0.61 |

Table S3: TWIST individual parameter sets and associated performance metrics for all acceptable solutions ($n = 17$), defined as parameter sets yielding calibration performance within 5% of the best-fit solution (see Methods). Reported are model parameters, objective function values (obj), normalization factor (q99), and performance metrics for calibration (c) and evaluation (e) periods at hourly (h) and daily (d) resolution.

| **ID** | $F_E$ | $F_{\text{TWD}}$ | $F_\theta$ | obj | q99 | cRMSE$_h$ | cRMSE$_d$ | cBias$_h$ | cBias$_d$ | c$R_h^2$ | c$R_d^2$ | eRMSE$_h$ | eRMSE$_d$ | eBias$_h$ | eBias$_d$ | e$R_h^2$ | e$R_d^2$ |
|---|---|---|---|---|---|---|---|---|---|---|---|---|---|---|---|---|---|
| 1 | 0.83 | 0.18 | 0.84 | 0.88 | 0.99 | 0.23 | 0.19 | -0.02 | -0.02 | 0.92 | 0.94 | 0.41 | 0.40 | -0.03 | -0.03 | 0.42 | 0.42 |
| 2 | 0.84 | 0.19 | 0.86 | 0.88 | 0.99 | 0.23 | 0.19 | -0.01 | -0.01 | 0.92 | 0.94 | 0.41 | 0.40 | -0.02 | -0.02 | 0.42 | 0.42 |
| 3 | 0.88 | 0.19 | 0.84 | 0.88 | 0.94 | 0.23 | 0.19 | -0.04 | -0.04 | 0.92 | 0.94 | 0.41 | 0.40 | -0.05 | -0.05 | 0.42 | 0.42 |
| 4 | 0.84 | 0.18 | 0.78 | 0.88 | 0.93 | 0.23 | 0.19 | -0.04 | -0.04 | 0.92 | 0.94 | 0.41 | 0.40 | -0.05 | -0.05 | 0.42 | 0.42 |
| 5 | 0.83 | 0.18 | 0.76 | 0.89 | 0.92 | 0.23 | 0.19 | -0.04 | -0.04 | 0.92 | 0.94 | 0.41 | 0.40 | -0.05 | -0.05 | 0.42 | 0.42 |
| 6 | 0.85 | 0.20 | 0.95 | 0.89 | 1.03 | 0.23 | 0.19 | 0.01 | 0.01 | 0.92 | 0.94 | 0.42 | 0.40 | 0.00 | 0.00 | 0.42 | 0.42 |
| 7 | 0.91 | 0.21 | 0.93 | 0.89 | 0.99 | 0.23 | 0.19 | -0.04 | -0.04 | 0.92 | 0.94 | 0.41 | 0.39 | -0.04 | -0.04 | 0.42 | 0.42 |
| 8 | 0.74 | 0.19 | 0.85 | 0.89 | 1.03 | 0.23 | 0.19 | 0.00 | 0.00 | 0.92 | 0.94 | 0.41 | 0.39 | 0.00 | 0.00 | 0.42 | 0.42 |
| 9 | 0.78 | 0.20 | 0.90 | 0.89 | 1.03 | 0.23 | 0.19 | -0.01 | -0.01 | 0.92 | 0.94 | 0.41 | 0.39 | -0.01 | -0.01 | 0.42 | 0.42 |
| 10 | 0.87 | 0.20 | 0.88 | 0.89 | 0.98 | 0.23 | 0.19 | -0.04 | -0.04 | 0.92 | 0.94 | 0.41 | 0.39 | -0.04 | -0.04 | 0.42 | 0.42 |
| 11 | 0.72 | 0.20 | 0.89 | 0.90 | 1.04 | 0.23 | 0.20 | -0.01 | -0.01 | 0.92 | 0.94 | 0.40 | 0.39 | 0.00 | 0.00 | 0.42 | 0.42 |
| 12 | 0.74 | 0.17 | 0.81 | 0.91 | 1.03 | 0.23 | 0.20 | 0.03 | 0.03 | 0.92 | 0.94 | 0.42 | 0.41 | 0.01 | 0.01 | 0.42 | 0.42 |
| 13 | 0.62 | 0.17 | 0.79 | 0.91 | 1.06 | 0.24 | 0.20 | 0.02 | 0.02 | 0.91 | 0.94 | 0.41 | 0.40 | 0.02 | 0.02 | 0.42 | 0.42 |
| 14 | 0.81 | 0.16 | 0.69 | 0.91 | 0.89 | 0.23 | 0.20 | -0.02 | -0.02 | 0.92 | 0.94 | 0.42 | 0.41 | -0.04 | -0.04 | 0.42 | 0.42 |
| 15 | 0.71 | 0.18 | 0.76 | 0.91 | 1.00 | 0.23 | 0.20 | -0.04 | -0.04 | 0.92 | 0.94 | 0.40 | 0.38 | -0.03 | -0.03 | 0.42 | 0.42 |
| 16 | 0.87 | 0.20 | 0.96 | 0.91 | 1.04 | 0.24 | 0.20 | 0.03 | 0.03 | 0.92 | 0.94 | 0.42 | 0.41 | 0.02 | 0.02 | 0.42 | 0.42 |
| 17 | 0.89 | 0.19 | 0.77 | 0.92 | 0.89 | 0.23 | 0.20 | -0.07 | -0.07 | 0.92 | 0.94 | 0.41 | 0.40 | -0.08 | -0.08 | 0.42 | 0.42 |

### References


Peters, Richard L., David Basler, Roman Zweifel, David N. Steger, Tobias Zhorzel, Cedric Zahnd, et al. (2025). "Normalized tree water deficit: an automated dendrometer signal to quantify drought stress in trees". en. In: *New Phytologist*, nph.70266. DOI: 10.1111/nph.70266.

Ziegler, Yanick, Rüdiger Grote, Franklin Alongi, Timo Knüver, and Nadine K. Ruehr (2024). "Capturing drought stress signals: The potential of dendrometers for monitoring tree water status." In: *Tree Physiology*. S2ID: 71a5eabeabb5bf1957c5afc635f2cc11cba6c188. DOI: 10.1093/treephys/tpae140.
